# Mitochondrial Fission Regulates Transcription of Ribosomal Protein Genes in Embryonic Hearts

**DOI:** 10.1101/2021.02.10.430627

**Authors:** Qiancong Zhao, Shun Yan, Jin Lu, Danitra J. Parker, Huiying Wu, Qianchuang Sun, David K. Crossman, Shanrun Liu, Qin Wang, Hiromi Sesaki, Kasturi Mitra, Kexiang Liu, Kai Jiao

## Abstract

Mitochondrial dysfunction causes severe congenital heart diseases and prenatal/neonatal lethality. The lack of sufficient knowledge regarding how mitochondrial abnormalities affect cardiogenesis poses a major barrier for the development of clinical applications that target inborn heart defects due to mitochondrial deficiency. Mitochondrial morphology, which is regulated by fission and fusion, plays key roles in determining mitochondrial activity. *Drp1* encodes a dynamin-related GTPase required for mitochondrial fission. To investigate the role of mitochondrial fission on cardiogenesis during the embryonic metabolic shift period, we specifically inactivated *Drp1* in second heart field derived structures. Deletion of *Drp1* in embryonic cardiomyocytes led to severe defects in mitochondrial morphology, ultrastructure, and activity. These defects caused increased cell death, decreased cell survival, disorganized cardiomyocytes, and embryonic lethality. Through characterizing this model, we reveal a novel AMPK-SIRT7-GABPB axis that relays the mitochondrial fission anomaly to reduced transcription of ribosomal protein genes in mutant cardiomyocytes. We therefore provide the first mouse genetic evidence to show that mitochondrial fission is essential for embryonic heart development. Furthermore, we uncovered a novel signaling cascade that mediates the crosstalk between mitochondrial dysfunction and protein synthesis. Our research provides further mechanistic insight regarding how mitochondrial dysfunction causes pathological molecular and cellular alterations during cardiogenesis.

## Introduction

Originating from engulfment of α-proteobacteria by precursors of modern eukaryotic cells, mitochondria have now evolved to become a vital organelle with multiple critical functions in eukaryotes, including supplying energy, regulating apoptosis, maintaining calcium homeostasis, sustaining redox homeostasis, and generating critical metabolites for cellular activities (1–6). In addition to their roles in postnatal organs under normoxia, accumulated evidence has established that mitochondria are also critical for fetal organogenesis including embryonic heart development in hypoxia (3, 5). Mitochondrial dysfunction in fetal hearts may lead to severe congenital heart defects (CHDs) and prenatal/neonatal lethality in human patients (7–9). These heart defects cover a large spectrum including hypertrophic, dilated, and noncompaction cardiomyopathies, suggesting the complex regulatory role of mitochondria during heart development. To date, no specific treatment has been developed to target CHDs caused by mitochondrial dysfunction. This clinical deficiency is at least partially due to the lack of sufficient knowledge on how mitochondrial defects affect normal heart development at the molecular, cellular and tissue levels.

The heart is the first functional organ formed in mammalian embryos (10–12). In mouse embryos prior to E11.5, cardiac mitochondria are highly immature with few cristae protruding into the mitochondrial matrix, and cardiomyocytes almost exclusively rely on anaerobic glycolysis to acquire ATP (3, 13, 14). By E13.5, cardiac mitochondria are functionally mature with their electron transportation and oxidative phosphorylation (OXPHOS) activities being indistinguishable from those in mitochondria of adult hearts (13–15). At this stage, the ultrastructure of cardiac mitochondria is very similar to the fully matured mitochondria with tubular cristae connected with the periphery, and embryonic cardiomyocytes acquire energy through both anaerobic glycolysis and OXPHOS (5, 13, 14, 16, 17). By the end of gestation, >50% of total ATP in hearts is generated through OXPHOS with the remaining generated through anaerobic glycolysis (5, 18). Therefore, cardiac mitochondria already provide a large portion of energy to support heart pumping at mid-to late- gestation stages, and the process of cardiac metabolic shift from glycolysis to OXPHOS starts in embryos.

In response to extra-/intra- cellular stimuli, mitochondria undergo fission and fusion, leading to a highly dynamic tubular network in cells. Mitochondrial morphology is a major determining factor for the normal functions of mitochondria; disruption of mitochondrial fission or fusion through genetic and pharmacological approaches leads to defects in numerous *in vitro* and *in vivo* systems (3–6, 19, 20). A group of cellular dynamin-related GTPases are pivotal for mitochondrial dynamics, with MFN1/MFN2 and DRP1 required for fusion and fission, respectively (3–6, 19, 20). The function of mitochondrial dynamics on cardiogenesis is initially demonstrated by studies using the *Nkx2.5-Cre* driver to simultaneously inactivate *Mfn1* and *Mfn2* (21, 22), which have partially overlapping functions in promoting mitochondrial fusion (23, 24). Double inactivation of *Mfn1/Mfn2* in embryonic cardiomyocytes aberrantly increases cytosolic calcium concentration and calcineurin activity, leading to enhanced Notch signaling to impair expression of multiple cardiomyocyte differentiation genes (21). DRP1 is essential for mitochondrial fission; it is recruited to the outer mitochondrial membrane by various adaptor proteins (including FIS1, MFF, MiD49, and MiD51), oligomerizes, and subsequently leads to mitochondrial constriction and eventual fission (6, 19). In previous studies, *Drp1* was inactivated in hearts using *Myh6-Cre* and *MCK*-*Cre* (25, 26). While these studies have demonstrated that mitochondrial fission is critical for postnatal heart development, they did not address the role of mitochondrial fission during embryonic heart development, as expression of *Drp1* was not efficiently inactivated in fetal hearts in these studies.

In our current study, we aimed to test the effect of blocking mitochondrial fission on embryonic heart development during the crucial period when cardiac metabolic shift starts to occur. To this end, we specifically inactivated *Drp1* using the *Mef2c-AHF-Cre* driver (27). We show that deletion of *Drp1* in embryonic hearts impairs mitochondrial morphology, ultrastructure and function, leading to severe myocardial wall defects and embryonic lethality before E17.5. We thus provide the first mouse genetic evidence to support the essential role of *Drp1* during mammalian cardiogenesis. Furthermore, through characterizing the conditional knockout (cKO) hearts, we reveal that the mitochondrial dysfunction caused by *Drp1*-deficiency reduces transcription of a group of ribosomal protein (RP) genes through the AMPK-SIRT7-GABPB axis. While mitochondria dysfunction can repress mRNA translation through the integrated stress response (ISR) (28) or the mammalian target of rapamycin (mTOR) signaling pathway (29–31), our research has revealed a novel mechanism, i.e. repressing RP gene transcription, by which mitochondrial dysfunction downregulates protein synthesis. We thus provide further insight regarding how mitochondrial dysfunction results in pathological alterations during organogenesis in fetus.

## Results

### 1. Deletion of *Drp1* in embryonic hearts led to hypoplastic myocardial wall and embryonic lethality

To understand the function of *Drp1* during cardiogenesis, we used the *Mef2c-AHF-Cre* driver (27) to inactivate *Drp1* in second heart field (SHF) derived structures including the outflow tract (OFT) and right ventricle (RV). We crossed *Mef2c-AHF-Cre/Drp1^loxp/+^* male mice with *Drp1^loxp/loxp^* female mice (32) to obtain cKO (*Mef2c-AHF-Cre/Drp1^loxp/loxp^*) and control (*Drp1^loxp/+^* or *Drp1^loxp/loxp^*) embryos at various stages. We first tested at which stage expression of DRP1 was inactivated through immunostaining assays. At E9.5, DRP1 expression could be defected in ~50% of cardiomyocytes in mutant RVs (Sup. Fig. 1), and its expression was only observed in very few cardiomyocytes in the OFT and RV of mutant embryos at E10.5 (Fig. 1A), indicating that *Drp1* was efficiently inactivated in structures derived from SHF at this stage. Expression of DRP1 in the common atria and atrial-ventricular canal region remained unchanged in mutant hearts, confirming the specificity of this Cre line. Mutant embryos could be recovered at the Mendelian ratio (~25%) until E15.5. At E16.5, only 9% of total living embryos were mutants and no mutant embryo could be recovered at E17.5 (Fig. 1B), suggesting that embryonic lethality occurred between E15.5 and E17.5.

**Figure 1.**
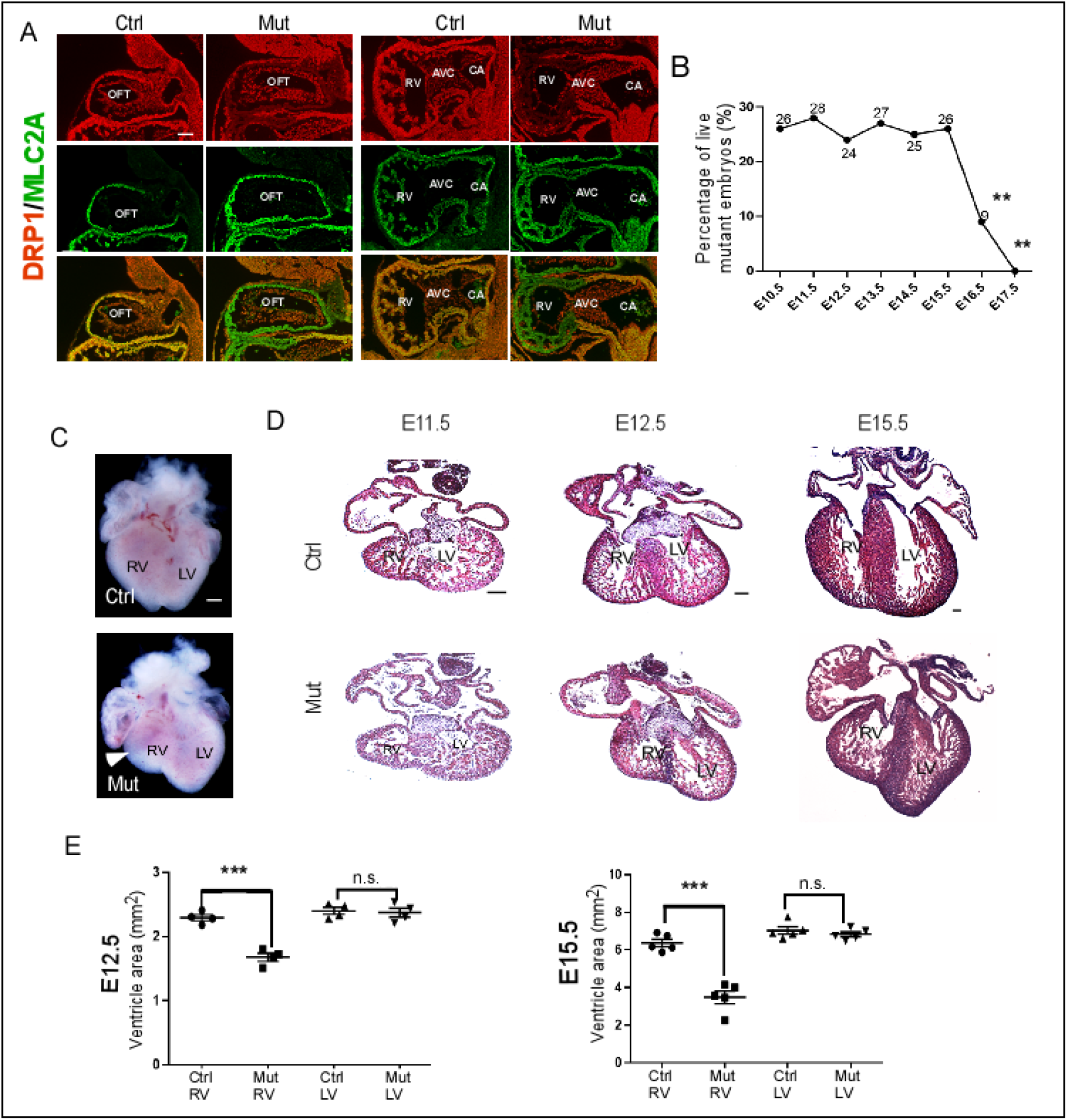
Deletion of *Drp1* by *Mef2c-AHF-Cre* led to severe defects in the right ventricle. *Mef2c-AHF-Cre/Drp1^loxp/+^* male mice were crossed with *Drp1^loxp/loxp^* female mice to obtain mutant (Mut, *Mef2c-AHF-Cre/Drp1^loxp/loxp^*) and control (Ctrl, *Drp1^loxp/+^* or *Drp1^loxp/loxp^*) embryos at different stages. **(A)** Sagittal sections of E10.5 embryos were immunostained with antibodies against DRP1 (red) and MLC2A (red, a cardiomyocyte marker). The outflow tract (OFT), atrial ventricular canal (AVC) and common atrium (CA) regions are shown. Expression of DRP1 was efficiently inactivated in the OFT and RV of mutant hearts but not other areas. **(B)** Percentage of live mutant embryos at different stages. Data were acquired from at least 4 litters at each developmental stage. **: *P<0.01*, Chi-square test. Embryonic lethality started to occur between E15.5 and E16.5. **(C)** Whole mount of control and mutant hearts at E14.5. The arrow indicates the reduced size of the mutant RV. **(D)** HE staining of heart sections at different stages. Reduced size of mutant RVs started to be observed from E12.5. At E15.5, mutant hearts also displayed non-compaction cardiomyopathy. **(D)** Quantification of the size of RVs and LVs in mutant and control hearts at E12.5 (left chart) and E15.5 (right chart). The size of RV and LV was measured using Photoshop 11. The area of each structure was calculated as the average of 5 consecutive sections with the largest area. ***: *P<0.001*, two tailed unpaired Student’s test; n.s.: not significant. Data are shown as mean ± SEM (Standard error). Scale bars in panels A, C and D represent 100um.

Whole mount examination showed that the size of the mutant RV was reduced at E14.5 and this hypoplastic RV phenotype was confirmed in embryos from E12.5 to E15.5 by histological studies (Fig. 1C-E). No overt defect was observed in mutant RVs at E11.5. The size of the left ventricle (LV) was not altered in all stages examined. In addition to the reduced size of RVs, we also observed the noncompaction defect at E15.5 in both RVs and LVs of mutant hearts (Fig. 1D). Since *Drp1* was specifically inactivated in SHF derived structures, the defect in the LV of mutants is thus secondary to the RV defect, supporting the idea that development of the two ventricular chambers is coordinated. The noncompaction defect observed in mutant hearts is consistent with the clinical observation that mitochondrial dysfunction is a primary cause of congenital noncompaction (33, 34). No defect was observed in the OFT region including aorta and pulmonary trunk. We thus show, for the first time, that DRP1 is required for normal cardiogenesis.

### 2. Mutant cardiomyocytes displayed abnormal cell proliferation, survival and orientation

To understand how deletion of *Drp1* leads to hypoplastic RVs, we examined cell proliferation and apoptosis. No abnormality was observed in mutant hearts at E11.5. Starting from E12.5, we observed significantly decreased cell proliferation and increased apoptosis in the RV of mutant hearts (Fig. 2). No cell proliferation or survival defect was observed in mutant LVs. During examination of cell proliferation and death, we noticed that cardiomyocytes in mutant RVs appeared to be less well organized than in control samples. We therefore examined expression of N-Cadherin, which mediates cell-cell interaction and is required for normal orientation of cardiomyocytes in mouse embryos (35, 36). Our N-Cadherin staining in Fig. 3A confirmed that mutant cardiomyocytes were less well organized in mutant RVs. In addition, the N-Cadherin immunostaining signal appeared be lower in cKO cardiomyocytes, and this result was confirmed through subsequent Western analysis (Fig. 3B). Reduced expression of N-Cadherin suggests abnormal formation of cell-cell contacts in mutant samples. We next directly examined the orientation of cardiomyocytes in the compact zone of LVs and RVs as described previously (35). At E12.5, the majority of cardiomyocytes (38/50) oriented perpendicularly to the heart wall in control RVs, while most cardiomyocytes in mutant RVs (40/50) were parallel to the heart wall (Fig. 3C, D). The orientation of cardiomyocytes in mutant LVs was not altered. Our data collectively indicate that *Drp1* is required for normal cell proliferation, survival and orientation in developing hearts.

**Figure 2.**
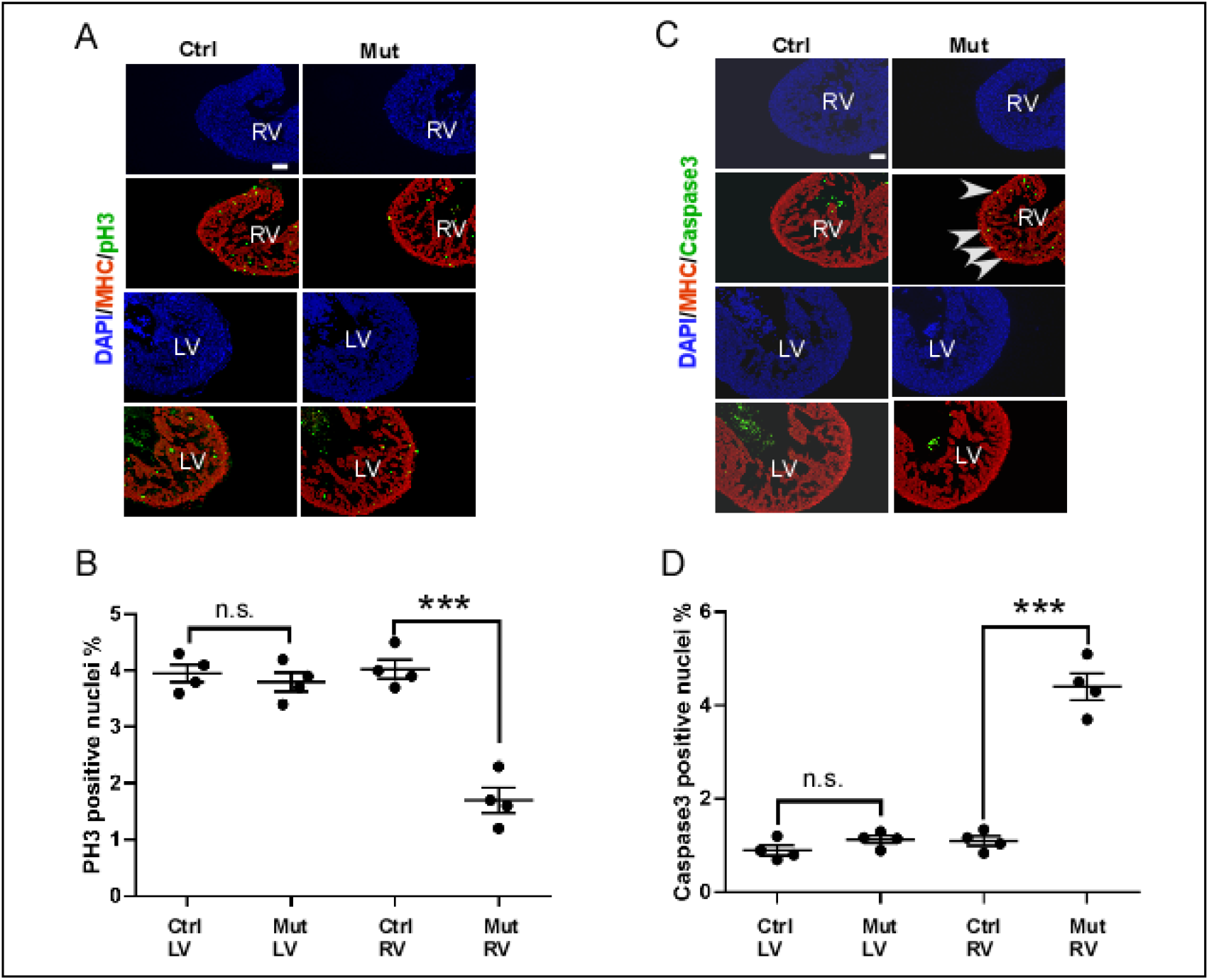
Deletion of *Drp1* led to abnormal cell proliferation and survival. **(A)** Sagittal sections of E12.5 hearts from mutant (Mut, *Mef2c-AHF-Cre/Drp1^loxp/loxp^*) and control (Ctrl, *Drp1^loxp/+^* or *Drp1^loxp/loxp^*) embryos were co-stained with an anti-phospho-histone 3 (pH3) antibody (green) and an anti-myosin heavy chain (MHC) antibody (red). Total nuclei were visualized with DAPI staining (blue). Examples of both left ventricles (LVs) and right ventricles (RVs) are shown. **(B)** Quantification results of pH3 staining. **(C)** Sagittal sections of E12.5 control and mutant hearts were co-stained with an anti-cleaved Caspase 3 antibody (green) and an anti-MHC antibody (red). Arrows show examples of positively stained nuclei. **(D)** Quantification results of Caspase 3 staining. ***: *P<0.001*, two tailed unpaired Student’s test; n.s.: not significant. Data are shown as mean ± SEM. Scale bars in panels A and C represent 50um.

**Figure 3.**
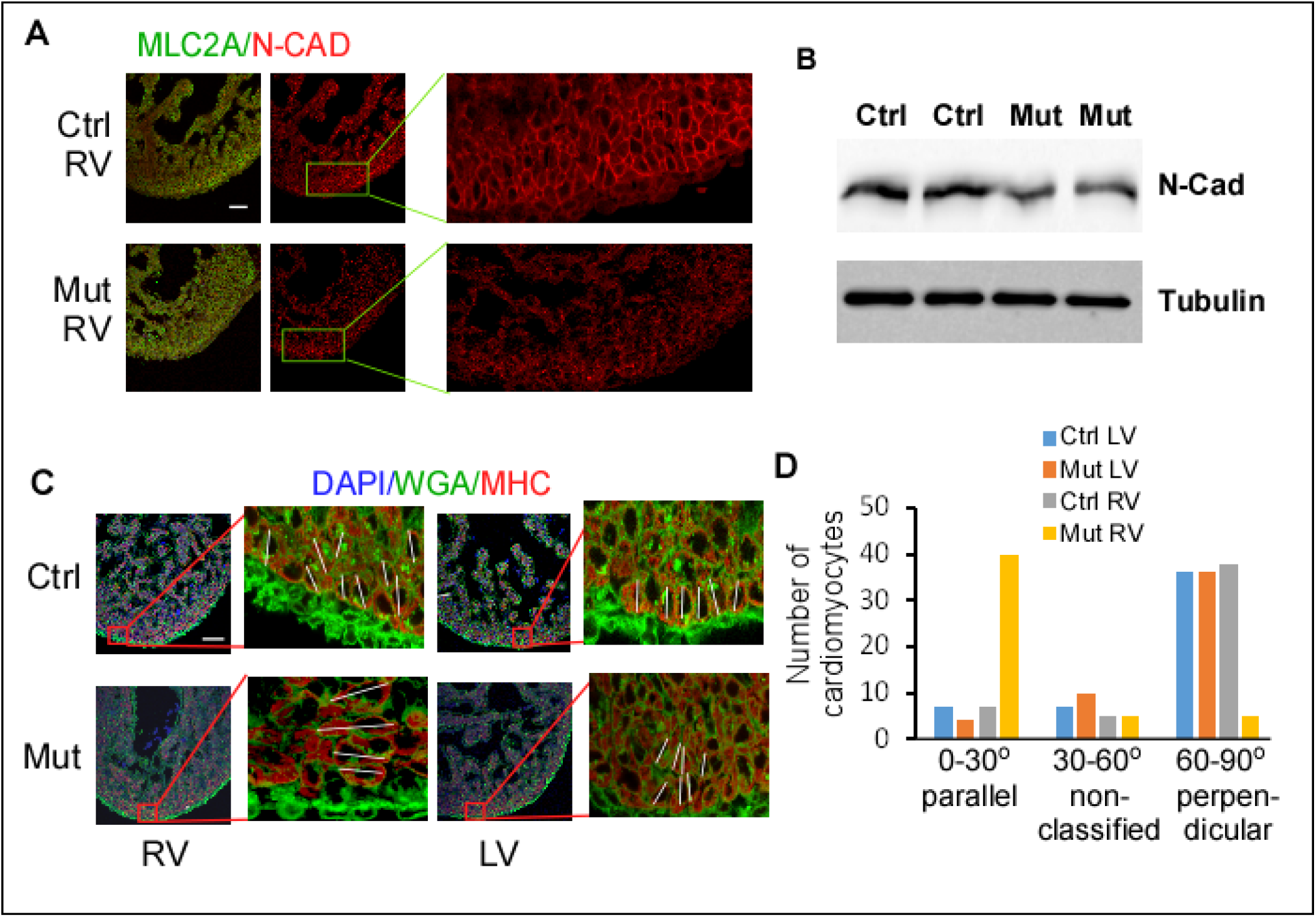
Deletion of *Drp1* led to abnormal cardiomyocyte orientation. **(A)** Sagittal sections of E12.5 hearts from mutant (Mut, *Mef2c-AHF-Cre/Drp1^loxp/loxp^*) and control (Ctrl, *Drp1^loxp/+^* or *Drp1^loxp/loxp^*) embryos were co-stained with an anti-N-Cadherin (N-Cad) antibody (green) and an anti-MLC2A antibody (red). **(B)** Expression of N-Cadherin in control and mutant RVs at E12.5 was examined through Western analysis. Tubulin was used as a loading control. **(C)** E12.5 heart sections were stained with WGA to label the cell membrane (green) and an anti-MHC antibody (red) to label cardiomyocytes. Nuclei were visualized with DAPI staining. We measured the orientation cardiomyocytes in the compact zone of control and mutant LVs and RVs following published procedures (35). A total of 50 cardiomyocytes for each structure (from 3 embryonic hearts) were measured. The white lines show the longitudinal axis of a cell. **(D)** The number of cardiomyocytes with different orientations relative to the myocardial wall (parallel, non-classified, and perpendicular). The distribution of cardiomyocytes of mutant RVs in the three categories was significantly different from that of the control LVs (*P<0.001*, Fisher’s exact test, “stats” package in R 4.0.3), while no significant difference was observed between control and mutant LVs. Scale bars in panels A and C represent 50um.

### 3. *Drp1*-deficiency caused defects in mitochondrial morphology and ultrastructure

To determine how deletion of *Drp1* affects mitochondrial morphology, we first examined cultured primary cardiomyocytes derived from cKO and control RVs at E10.5 (Fig. 4A). We co-stained cells with antibodies for a mitochondrial marker, TOM20 (37), and DRP1. In comparison to the control cells, cKO cells exhibited efficient reduction in DRP1 expression and displayed characteristic tubular mitochondria (Fig. 4A), as expected (20). To confirm this observation in embryonic cardiac tissues, we stained E10.5 heart sections with TOM20 and DRP1 antibodies. We found that mitochondria in many cKO cells were clustered at one side of nuclei in contrast to their even distribution in control cells (Fig. 4B). This mitochondrial clustering phenotype is similar to the morphology observed in Drosophila tissue with *Drp1* inactivated (38). Therefore, *Drp1*-deletion led to characteristic tubular mitochondrial morphology in developing cardiomyocytes.

**Figure 4.**
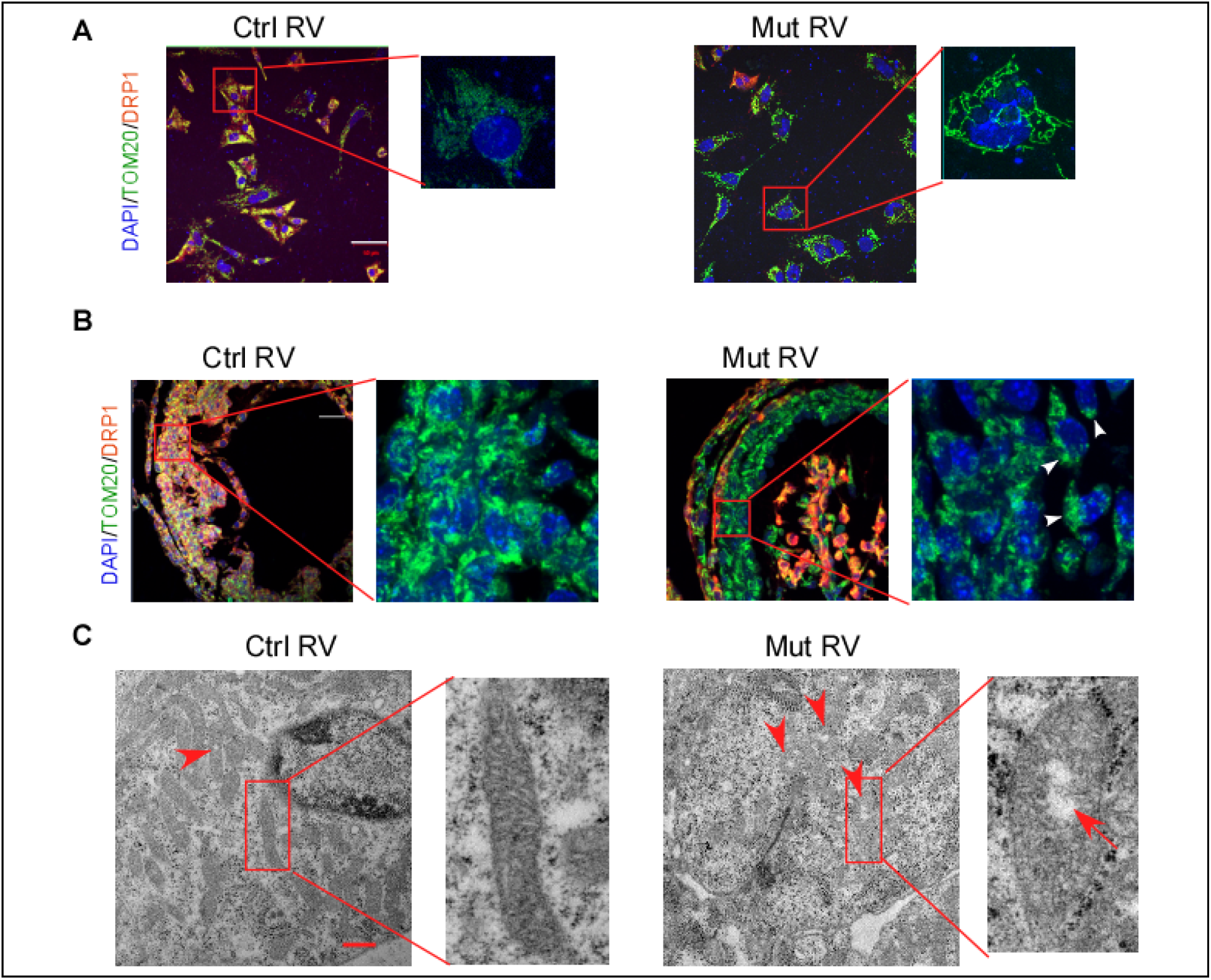
Deletion of *Drp1* impaired mitochondrial morphology and ultrastructure. **(A)** Cardiomyocytes were isolated from RVs of mutant (Mut, *Mef2c-AHF-Cre/Drp1^loxp/loxp^*) and control (Ctrl, *Drp1^loxp/+^* or *Drp1^loxp/loxp^*) embryos at E10.5 and were cultured for 48 hours. They were then co-stained with antibodies against TOM20 (green) and DRP1 (red). Nuclei were stained with DAPI (blue). Samples were examined using a confocal microscope. Mutant cells (without DRP1 expression) displayed tubular mitochondria. **(B)** Sagittal sections of E10.5 hearts were immunostained with antibodies against TOM20 (green) and DRP1 (red), and samples were examined using a confocal microscope. In many mutant cells, mitochondria were clustered to one side of the nucleus. The arrow heads shows some examples of such cells. **(C)** Sections of control and mutant embryonic hearts (E13.5) were examined through TEM. Arrows indicate examples of mitochondria that contain an area with reduced electron density in the matrix. Scale bars in panels A and B represent 50um, and the one in panel C represents 1um.

We next quantified mitochondrial DNA levels using quantitative polymerase chain reaction (qPCR). The result in Sup. Fig. 2A shows that the mitochondrial DNA in cKO RVs (E12.5) was not significantly different from control RVs. Consistent with this result, expression of TFAM, a key transcription factor for mitochondrial DNA transcription and replication (3), was not altered by *Drp1*-deficiency (Sup. Fig. 2B). This result is different from that obtained in adult cardiomyocytes, in which deletion of *Drp1* reduced TFAM expression and mitochondrial biogenesis (39). Our results suggest that the interaction between mitochondrial dynamics and biogenesis has not been fully established in embryonic cardiomyocytes.

We next examined mitochondrial ultrastructure using Transmission electron microscopy (TEM). Significantly, we observed that ~14% (85/611, from 3 biological replicates) of mitochondria of cKO cardiomyocytes (E13.5) contained an area with reduced electron density in their matrices (Fig. 4C). Whereas, <1% (6/647) of mitochondria from control cardiomyocytes showed a similar phenotype. In these mitochondria, their cristae appeared to be disorganized, similar to immature cardiac mitochondria. We thus showed that *Drp1*-deletion led to an abnormal ultrastructure of mitochondria in cardiomyocytes.

### 4. Deletion of *Drp1* impaired OXPHOS activity in embryonic cardiomyocytes

We speculated that abnormal morphology and ultrastructure would lead to reduction of the mitochondrial activity in mutant cardiomyocytes. We thus examined electron transportation chain (ETC) function by assessing oxygen consumption of RV tissues (E14.5) (Fig. 5A, B), and calculated mitochondrial respiration rate as previously described (15). There was no significant difference in basal respiration rate (BL) between control and cKO samples, nor when triggered by Complex I substrates (malate + glutamate) (V_0_) (Fig. 5A). However, the ADP induced maximum respiration rate (V_max_) was significantly lower in cKO samples than that in control samples (Fig. 5C), suggesting that the oxidation phosphorylation capacity of complex I was impaired by *Drp1*-deletion. The significant reduction in respiratory control ratio (RCR, V_max_/V_0_) with deletion of *Drp1* (Fig. 5B) suggests that the coupling of ETC with OXPHOS was compromised in cKO samples. In both cKO and control samples, addition of an ADP/ATP translocase inhibitor atractyloside (ATR) significantly reduced the mitochondrial respiration rate (Fig. 5A), indicating no detectable defect in ADP or ATP translocation across the inner membrane. To further test the effect of impaired ETC activity on generation of ATP, we measured the total ATP level in LVs and RVs at E13.5. As shown in Fig. 5C, the ATP level in cKO RVs was significantly reduced compared to the control level, while no significant difference was observed between control and cKO LVs.

**Figure 5.**
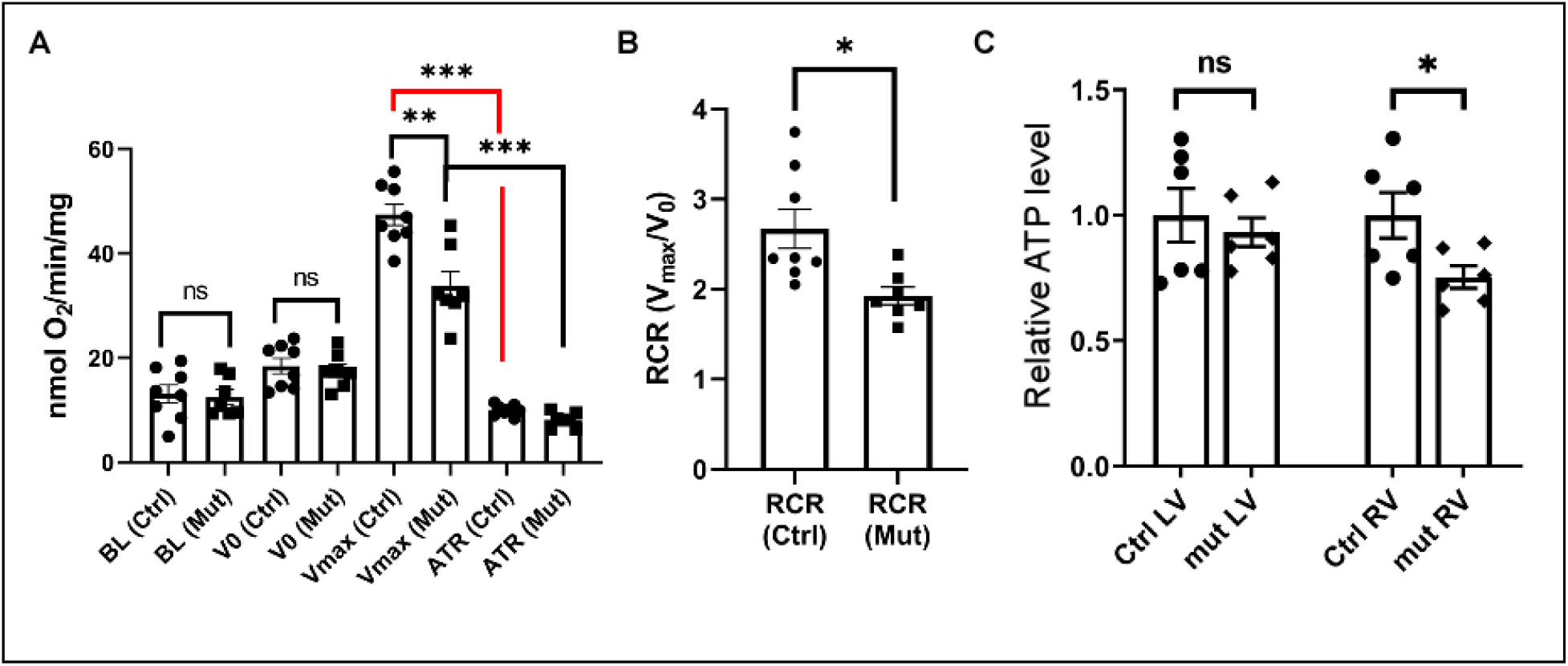
The effect of *Drp1*-deletion on OXPHOS in embryonic cardiomyocytes. **(A)** Oxygen consumption of E14.5 control and mutant RVs were examined using Oroboros. BL: base line. **(B)** RCR was calculated. **(C)** The ATP level in E13.5 LVs and RVs were measured. Data were normalized against the weight of samples. The level of control samples was set at 1.0. Data are shown as mean ± SEM. *: *P<0.05;* **: *P<0.01;* ***: *P<0.001*, two tailed unpaired Student’s t test.

### 5. Deletion of *Drp1* reduced transcription of multiple RP genes in embryonic cardiomyocytes

To better understand how deletion of *Drp1* affects cardiomyocyte development at the molecular level, we performed droplet-based single cell-RNA-Seq (scRNA-Seq) using cells isolated from control and cKO RVs at E13.5. A total of 7739 control cells and 6967 cKO cells passed the quality control and were used for data analysis. Using the Uniform Manifold Approximation and Projection (UMAP) dimension reduction technique (40), and based on known molecular markers of cells in embryonic hearts (41, 42), we identified cell populations corresponding to different cell types, including cardiomyocytes, endocardial cells, epicardial cells, endothelial cells, red blood cells, and macrophages (Fig. 6A, B, Sup. Fig. 3). To understand how deletion of *Drp1* specifically impact gene expression in developing cardiomyocytes, we combined the 3 cardiomyocyte clusters into one group and compared gene expression within this combined cluster between control and mutant cells. We found 207 genes whose expression was significantly upregulated or downregulated by at least 50% (adjusted *P*<0.05). We noticed that >10 ribosomal protein (RP) genes were among the most significantly downregulated genes (Fig. 6C, Sup. Fig. 4, Sup. Table 1). We then performed gene ontology (GO) term enrichment analysis using Metascape (43) (Fig. 6D, Sup. Table 2). The term “eukaryotic translation initiation”, which includes multiple RP genes, was most significantly enriched in the list, consistent with our observation in Fig. 6C. As expected, multiple pathways related to mitochondrial activities, such as OXPHOS, Respiratory chain complex I, and cytochrome c oxidase (complex IV), are impaired by deletion of *Drp1* (Fig. 6D). In addition, genes involved in muscle structure development (including *Tnnt1*, *Tnnt2*, *Actc1*, etc.) were also enriched in the list, suggesting that differentiation of cardiomyocytes was also compromised by *Drp1*-deletion. We did not observe altered Notch signaling in contrast to *Mfn1/2* double knockout cardiomyocytes (21), suggesting that the molecular defects caused by mitochondrial fission deficiency is different from that of mitochondrial fusion deficiency.

**Fig. 6.**
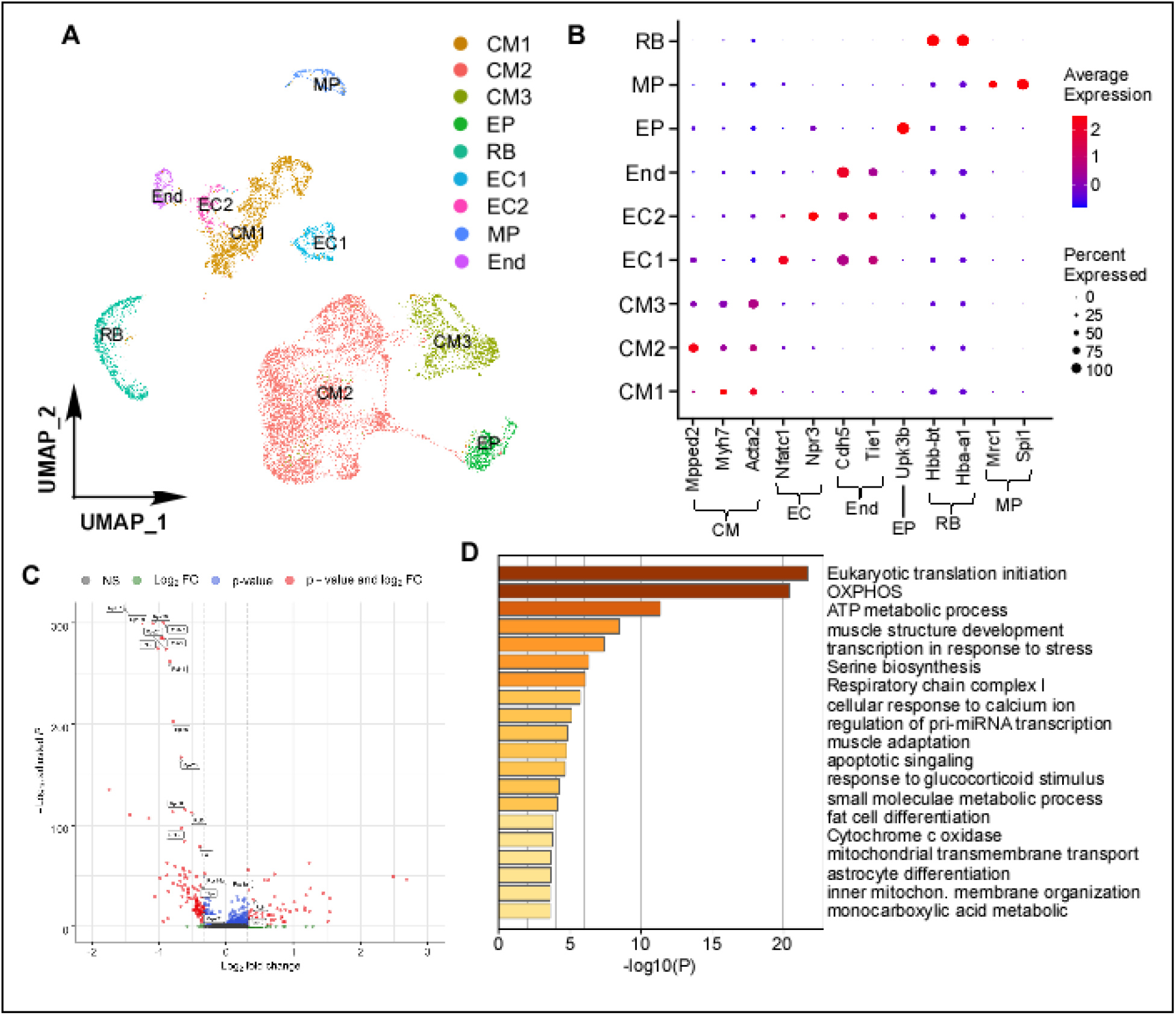
Expression of a group of RP genes in embryonic cardiomyocytes was impaired by *Drp1*-deletion. **(A)** ScRNA-Seq was performed using cells isolated from RVs of control and mutant embryos at E13.5. This chart shows UMAP projection of various cell types including cardiomyocytes (CM1, 2, 3), epicardial cells (EP), red blood cells (RB), endocardial cells (EC1, 2), macrophages (MP), and endothelial cells (End). **(B)** The Dot plot chart shows average expression of representative marker genes of different cell types across all clusters. **(C)** All three CM clusters were combined, and 207 genes were identified that were differentially expressed between control and mutant cardiomyocytes by at least 50% with an adjusted *P<0.05* (the red dots). The RP genes were indicated. The X-axis shows the folds of alteration and the Y-axis shows the adjusted P value. The dotted lines shows the threshold (adjusted *P<*0.05, expression altered by at least 50%). The same chart with a higher magnitude was provided in Sup. Fig. 4. The 207 genes underwent GO term enrichment analysis using Metascape. The top 20 terms are shown.

### Deletion of *Drp1* caused reduction in *de novo* protein synthesis in cKO cardiomyocytes

Few studies have made the connection between mitochondrial dysfunction and altered transcription of RP genes, and therefore we decided to focus on this group of genes. We first examined the expression of 10 RP genes through quantitative reverse transcription PCR (qRT-PCR) using RNA samples isolated from control and cKO RVs at E13.5. Our result in Fig. 7A confirmed reduced expression of these genes in cKO samples. Our Western analysis confirmed reduced expression of 4 RPs in mutant RVs at the protein level (Fig. 7B). To test the impact of reduced expression of RP genes, we examined protein synthesis in cultured cardiomyocytes derived from control and mutant RVs at E13.5. Our results show that protein synthesis was significantly reduced in cKO cardiomyocytes (Fig. 7C, D), as expected from the reduced expression of multiple RP genes.

**Fig. 7.**
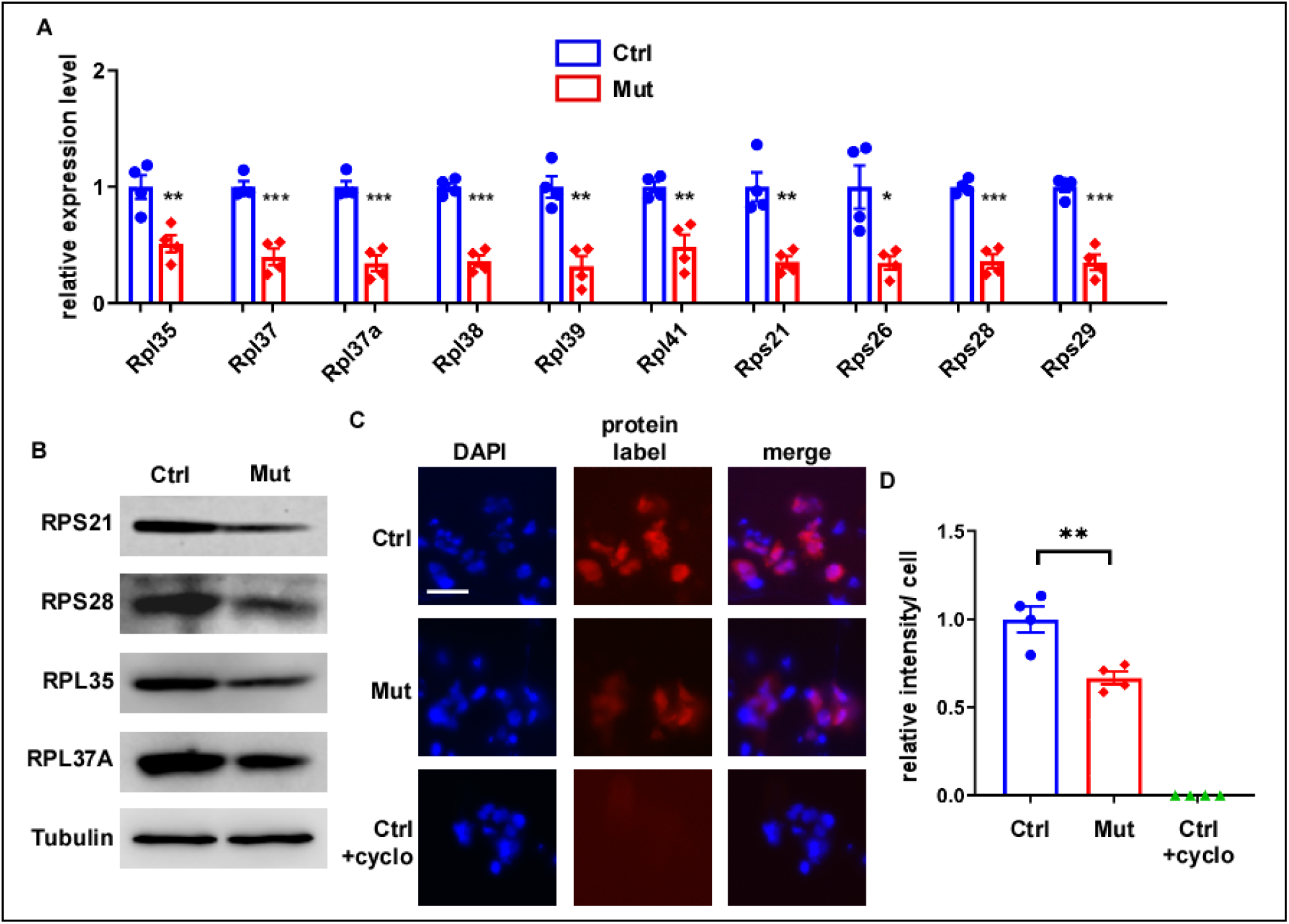
De novo protein synthesis was impaired by deletion of *Drp1*. **(A)** Total RNA were isolated from RVs of mutant (Mut, *Mef2c-AHF-Cre/Drp1^loxp/loxp^*) and control (Ctrl, *Drp1^loxp/+^* or *Drp1^loxp/loxp^*) hearts at E13.5 followed by qRT-PCR analysis to examine expression of multiple RP genes. Data were normalized against *Hprt*. The level of control samples was set at 1.0. Consistent with the scRNA-Seq data in Fig. 6, expression of multiple RP genes was significantly reduced by *Drp1*-deletion. **(B)** Proteins were isolated the RVs of control and mutant samples at E13.5. Western analysis was then performed using indicated antibodies. Tubulin was used as a loading control. **(C)** De novo protein synthesis was examined on cultured cardiomyocytes isolated from E13.5 hearts. Nuclei were visualized with DAPI staining (blue). The red signal labeled newly synthesized proteins. Control cells treated with Cycloheximide (cyclo, a protein synthesis blocker) was included as a negative control. **(D)** Quantification of protein synthesis. We measured the signal intensity of >100 cells for each culture. The total intensity was divided by the number of cells to obtain intensity/cell. The number of the control ones was set at 1.0. *: *P<0.05;* **: *P<0.01;* ***: *P<0.001*, two tailed unpaired Student’s t test.

### 7. Mitochondrial dysfunction activates the AMPK/SIR7/GABPB axis to reduce transcription of RP genes in cKO cardiomyocytes

In the next set of experiments, we aimed to determine the mechanism by which mitochondrial dysfunction leads to reduced transcription of RP genes. SP1, YY1 and GABP are major transcription factors promoting RP gene transcription (44–46); however, our initial Western analysis did not reveal reduced expression of these proteins in cKO samples (Fig. 8A). It is known that GABP is composed of two subunits, GABPA and GABPB, which contain the Ets DNA binding domain and transcriptional activation domain, respectively (47). We then performed a co-immunoprecipitation (co-IP) study to test formation of the transcriptional active GABP complex in mutant tissues. Data in Fig. 8B showed that the formation of the GABPA-GABPB complex was reduced by *Drp1*-deletion. It has been well established that post-translational modification of GABPB, including phosphorylation and acetylation, can interfere the interaction between GABPA and GABPB (48–50). We therefore re-probed the IP’ed samples with an antibody against phospho-Serine or against acetyl-Lysine. While the level of phospho-GABPB1 was not altered, the level of acetyl-GABPB1 was clearly increased (Fig. 8B), supporting the idea that increased acetylation of GABPB1 blocks its interaction with GABPA to form the active GABP complex.

**Fig. 8.**
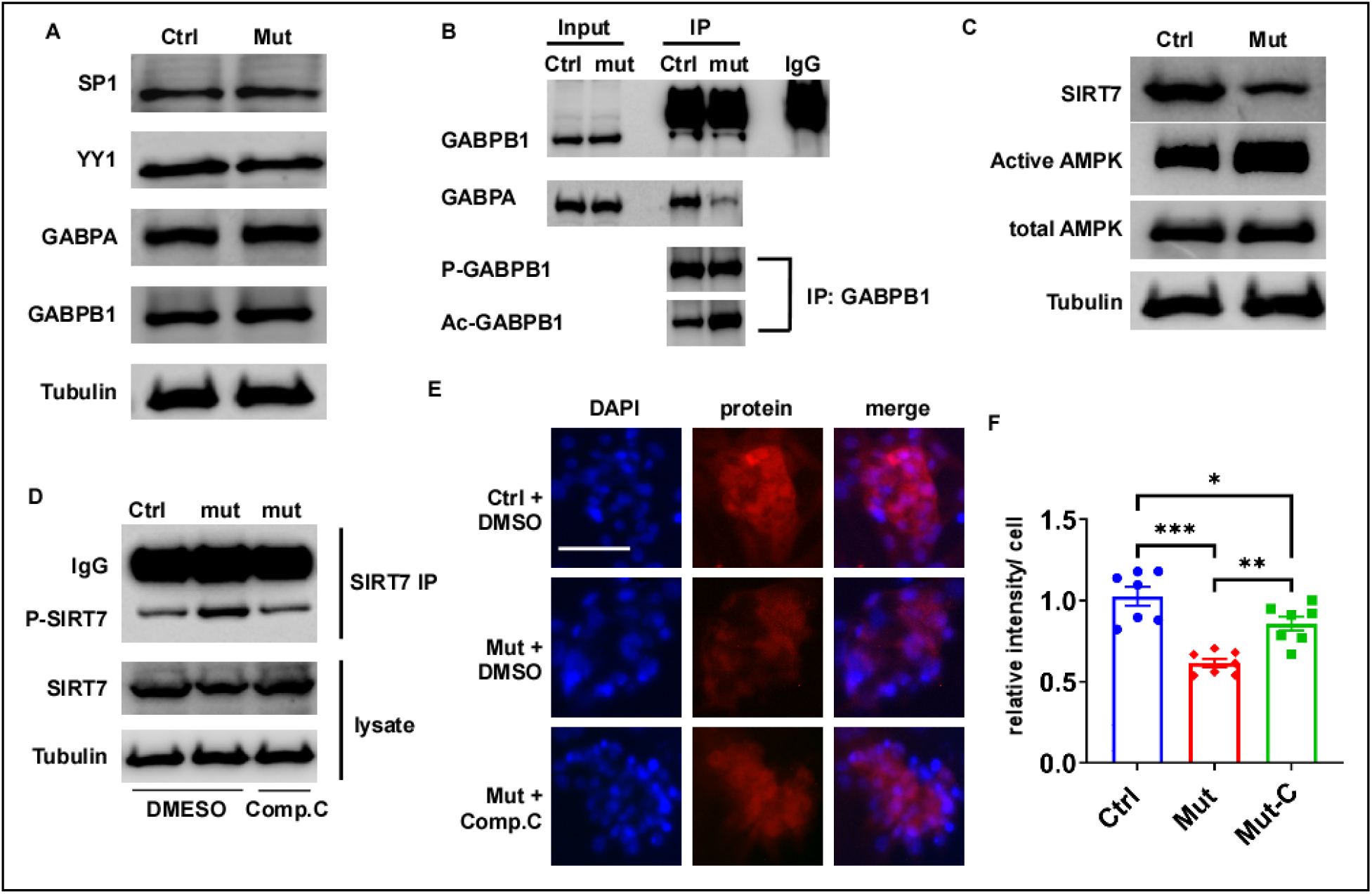
Deletion of *Drp1* led to reduced protein synthesis through the AMPK-SIRT7-GABPB axis. **(A)** Western analysis was performed on protein lysates isolated from RVs of mutant (Mut, Mef2c-AHF-Cre/Drp1loxp/loxp) and control (Ctrl, Drp1loxp/+ or Drp1loxp/loxp) embryos (E13.5) using indicated antibodies. Tubulin was used as a loading control. **(B)** GABPB1 was IP’ed from E13.5 RVs using an anti-GABPB1 antibody followed by Western analysis using various antibodies. Association of GABPA with GABPB1 was reduced in mutant samples. IP’ed samples were re-probed with an anti-phospho-Serine antibody or an anti-acetyl-Lysine antibody. The level of acetylated GABPB1 (Ac-GABPB1) was increased in mutant samples. **(C)** Western analysis was performed using E13.5 RV lysates. The level of SIRT7 was reduced in mutant samples. Total AMKP level remained the same while active AMKP was increased by *Drp1*-deletion. **(D)** E13.5 hearts were isolated, cultured, and treated with DMSO (control) or Compound C (Comp.C) for 6 hours. SIRT7 was IP’ed from the lysates of RVs followed by Western analysis using an anti-phospho-Serine antibody. Compound C treatment reduced the level of phosopho-SIRT7 (P-SIRT7) to the control level. Western analysis on total lysates showed that the total SIRT7 level was restored to the normal level in mutant samples by Compound C treatment. **(E, F)** Protein synthesis was measured in cultured cardiomyocytes treated with DMSO or Compound C. Quantification results in panel F show that Compound C treatment (Mut-C) partially rescued the protein synthesis defect in mutant samples. The scale bar represents 50um. *: *P<0.05;* **: *P<0.01;* ***: *P<0.001*, one way Anova followed by Tukey’s multiple comparisons test.

To further reveal how acetylation of GABPB1 was increased in cKO samples, we examined expression of SIRT7, which is a deacetylase of GABPB1 (50). As shown in Fig. 8C, expression of SIRT7 was downregulated by deletion of *Drp1*. Since the level of *Sirt7* mRNA was not altered in mutant cells from our scRNA-Seq data (Sup. Table 1), we examined whether phosphorylation of SIRT7 was altered, as this post-translational modification is known to alter its protein level (51). Our data in Fig. 8D clearly showed phosphorylation of SIRT7 was increased in cKO samples. Since AMP-activated protein kinase (AMPK) can directly phosphorylate SIRT7 upon energy starvation (51), we examined the active level of AMPK in cKO RVs. We show that active AMPK was increased by *Drp1*-deletion (Fig. 8C), and furthermore the level of phospho-SIRT7 in cKO RVs was restored to the control level by treatment of Compound C (an AMPK inhibitor). These data support the role of AMPK on SIRT7 phosphorylation. Finally, we showed that blocking AMPK through treatment of cKO cardiomyocytes with Compound C could partially rescue the global protein synthesis defect caused by *Drp1*-deletion (Fig. 8E, F), providing direct evidence to support the role of the AMPK-SIRT7 in mediating the protein synthesis defect in cKO cells (also see the model in Fig. 9).

**Fig. 9.**
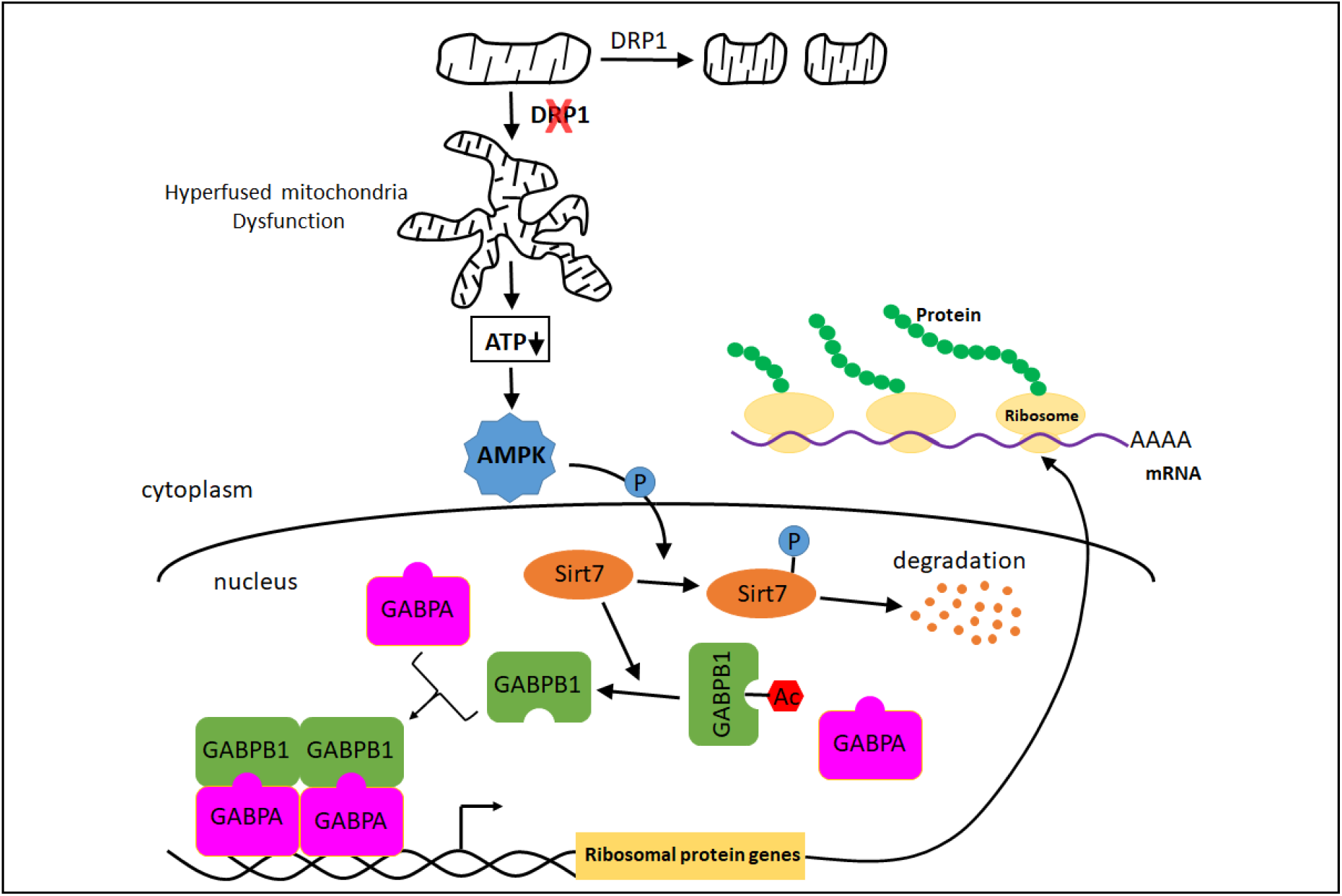
Model for how the defect in mitochondrial fission leads to reduced protein synthesis through repressing RP gene transcription.

## Discussion

To better understand the mechanism by which mitochondria regulate cardiogenesis, we have examined how blocking mitochondria fission affects mammalian cardiogenesis during the critical embryonic metabolic shift period in this study. In mouse embryos between E11.5 and E13.5 cardiac mitochondria undergo a maturation process enabling cardiomyocytes to obtain ATP from both anaerobic glycolysis and OXPHOS (5, 13, 14, 16, 17). We speculated that proper mitochondrial morphology would be important for mitochondrial maturation to support the embryonic metabolic shift, and therefore applied a conditional gene inactivation approach to delete *Drp1* in developing hearts.

Inactivation of *Drp1* in neonatal hearts results in animal lethality within the first 2 weeks after birth (25, 26), indicating that mitochondrial fission is essential for postnatal heart homeostasis. However, the effect of *Drp1-*deletion on embryonic heart development has not been reported in the literature. We inactivated *Drp1* using the *Mef2c-AHF-Cre* driver, which starts to inactivate target genes in the precursor cells of the SHF (27). Strong reduction in DRP1 expression was observed starting from E9.5 (Sup. Fig. 1), and by E10.5, DRP1 was only detected in few cardiomyocytes in the OFT and RV regions (Fig. 1). No reduction in DRP1 expression was observed in LVs of cKO hearts, and therefore cardiomyocytes in LVs of mutant hearts can be used as an internal control. We thus provide an ideal model to test the role of *Drp1* on cardiogenesis during cardiac mitochondrial maturation. Our immunofluorescence microscopy studies confirmed that mitochondria in mutant cardiomyocytes displayed signature fission defects (Fig. 4), including tubular mitochondria observed in cultured cells (Fig. 4A) and uneven clustering observed on tissue sections (Fig. 4B). Our following examination of mitochondrial ultrastructure through TEM showed that ~14% of mitochondria in cKO cardiomyocytes contained an area with reduced electron density (Fig. 5). The cristae in this type of mitochondria are disorganized, resembling immature mitochondria. Our data strongly suggest that mitochondrial fission is essential for proper mitochondrial maturation in developing hearts. Our functional tests have provided further support for the idea that abnormal mitochondrial morphology can lead to reduced mitochondrial activities (Fig. 5). Our results are in contrast to proliferating mouse embryonic fibroblasts or T cells, in which *Drp1* ablation can sustain elevated respiration (52–55). Therefore, the effect of *Drp1*-deletion on mitochondrial activities is influenced by cell types.

No defect in cKO cardiomyocytes was observed until E11.5. Starting from E12.5, *Drp1*-deletion in cardiomyocytes led to reduced cell proliferation, increased cell death and abnormal cell orientation (Fig. 1–3). This timeline correlates well with the mitochondrial maturation process; the cellular defects start to present when mitochondria begin to generate ATP through OXPHOS. We therefore provide another piece of evidence to support the idea that cardiac metabolic shift has already begun at mid-gestation and this shift is important for normal fetal heart development. Abnormal orientation of cardiomyocytes is a new phenotype associated with *Drp1*-deletion. Expression of N-Cadherin in *Drp1*-deleted cardiomyocytes was clearly reduced as shown from both immunostaining and Western analysis (Fig. 3). Considering the result from a recent publication that N-Cadherin is required for the proper orientation of cardiomyocytes (35), we speculate that DRP1-mediated mitochondrial fission acts through N-Cadherin to promote normal cell orientation. Our scRNA-Seq data failed to reveal reduced expression of N-Cadherin at the RNA level in mutant samples, meaning that modulation of N-Cadherin by *Drp1*-deletion likely occurs through certain post-transcriptional regulatory mechanisms. This result is different from the observation made in P19 teratocarcinoma cells; in these cells, transcription of N-Cadherin could be repressed by altering DRP1 expression (56). Thus, the mechanisms underlying the crosstalk between DRP1 and expression of N-Cadherin varies depending on cell types. Interestingly, expression of N-Cadherin could also be reduced by blocking mitochondrial fusion in mammary epithelial cells, the counter process of fission (57). Therefore, expression of N-Cadherin appears to be commonly targeted in different cell types by altered mitochondrial morphology.

One of the most significant discoveries of our current study is to reveal the AMPK-SIRT7-GABP axis that relays the mitochondrial fission defect to reduced cytosolic protein translation in mutant cells. The model shown in Fig. 9 summarizes our major discoveries. *Drp1*-deletion in embryonic cardiomyocytes blocks mitochondrial fission, leading to abnormal mitochondrial dynamics and reduced capacity to generate ATP. The reduced level of ATP activates AMPK, a key sensor of the energy level in cells (58), which phosphorylates SIRT7, leading to SIRT7 degradation. The normal function of SIRT7 is to remove the acetyl group from GABPB1 to promote the formation of the GABPA and GABPB complex, which activates transcription of multiple RP genes. The reduced level of SIRT7 due to increased phosphorylation by AMPK reduces RP gene transcription and represses cytosolic protein translation in cKO cardiomyocytes.

Two major pathways have been reported to mediate the crosstalk between mitochondrial stress and protein synthesis in mammalian cells. Mitochondrial dysfunction can act through the OMA1-DELE1-HER1 cascade to trigger the integrated stress response (ISR) (28), which represses global mRNA translation and at the same time enhances translation of some specific mRNAs including *ATF4*, *ATF5* and *DDIT3* (31). These proteins activate transcription of cytoprotective genes to help restore the function of mitochondria (31, 59, 60). In another pathway, the reduced energy level in cells due to mitochondrial dysfunction represses the activity of the mammalian target of rapamycin (mTOR) signaling pathway to inhibit protein synthesis (29–31). To the best of our knowledge, our research provides the first example showing that mitochondrial dysfunction can act through downregulating transcription of RP genes to repress protein synthesis. It will be of great interest to determine in future studies whether this signaling cascade can also function in other cell types. We noticed that blocking AMPK only partially rescued the protein synthesis defect in mutant cardiomyocytes (Fig. 8), suggesting the presence of another pathway, such as the ISR pathway, which acts independently with the AMPK-SIRT7-GABPB cascade to repress protein synthesis. Consistent with this idea, the pathway involved in transcription in response to stress was activated by *Drp1*-deletion (Fig. 6D).

A connection between *Drp1* and RP genes was also reported previously, although converse to what we report here (61, 62). Particularly, higher *Drp1* expression was found to correlate with lower expression of various ribosomal genes in tumor tissues of various cancer types (61). Similarly, a population of quiescent hematopoietic stem cells with higher DRP activity was found to be enriched in negative regulators of translation and RNA processing (62). Therefore, mitochondrial fission may increase or decrease RP gene transcription in a context dependent fashion.

In summary, we provide convincing mouse genetic evidence to show that *Drp1*-mediated mitochondrial fission is essential for normal development of embryonic cardiomyocytes. We further revealed a novel AMPK-SIRT7-GABPB signaling pathway that relay the mitochondrial fission defect to reduced RP gene transcription and protein synthesis. Our study helps us better understand the role of mitochondria in regulating cardiogenesis, and will contribute to development of novel clinical applications aimed at CHDs due to mitochondrial dysfunction.

## Methods

### 1. Mouse strains and maintenance

Mouse studies conform the Guide for the Care and Use of Laboratory Animals. Establishment of *Mef2c-AHF-Cre* mice (27) and *Drp1^loxp/loxp^* mice (32) and their genotyping strategy were described previously. *Mef2c-AHF-Cre* mice were maintained on the C57BL/6 and 129S6 mixed background, while *Drp1^loxp/loxp^* mice were maintained on the C57BL/6 background. *Mef2c-AHF-Cre* male mice were used to cross *Drp1^loxp/loxp^* mice to obtain *Mef2c-AHF-Cre*/*Drp1^loxp/loxp^* male mice, which were then crossed with female *Drp1^loxp/loxp^* mice to acquire control and cKO embryos at various stages. During mating, one male mouse and 1-2 female mice were put in the same cage, and the noon of detection of a plug in the vagina of a female mouse was designated as E0.5.

### 2. Tissue treatment, paraffin sectioning, HE staining, immunostaining and measuring cardiomyocyte orientation

Standard procedures were performed as described in previous studies (63–65). Briefly, embryos or embryonic hearts were dissected in PBS and fixed in 4% paraformaldehyde overnight at 4°C. The next day, samples were dehydrated with ethanol, cleared with Histoclear, and embedded in paraffin. Wax blocks were sectioned at 10um to acquire paraffin sections, which were kept at 4 °C until use. Sections were stained with Hematoxylin and Eosin (HE) for histological examination. For immunofluorescent staining, paraffin sections were dewaxed, rehydrated, treated with blocking buffer (10% serum in TBST (TBS with 0.1% Tween 20)), incubated with a primary antibody overnight at 4°C, and incubated with an Alexa-conjugated secondary antibody (ThermoFisher) at room temperature for 1 hour. Slides were then thoroughly washed with TBST and briefly stained with DAPI (ThermoFisher) to visualize the nuclei. If necessary, the TSA Plus Cyanine 3.5 System (PerkinElmer) was used to amplify the signal. Immunostained samples were observed using a Zeiss Axio fluorescent microscope or a Zeiss laser scanning confocal microscope (LSM700). The primary antibodies used for immunostaining were purchased from Abcam (DRP1, ab56788), BD Biosciences (N-Cadherin, #610920), Cell Signaling (cleaved Caspase3, #9661), Iowa Hybridoma Bank (MHC, MF20), Millipore (phospho-Histone H3, #06-570), Santa Cruz (TOM20, sc-11415). The anti-MLC2A antibody was kindly provided by Dr. S. Kubalak (Medical University of South Carolina). Measuring the orientation of cardiomyocytes in the compact zone was performed following the procedures described in a published study (35). E12.5 heart sections were stained with Wheat Germ Agglutinin (WGA, ThermoFisher) to label cell membrane as described previously (66). Sections were co-stained with an anti-MHC antibody to label cardiomyocytes. As previously defined (35), if the ratio of the length of a cell to its width is bigger than 120%, this cell is orientated, and its orientation was determined by the angle of the longitudinal axis to the myocardial wall surface reference line. Cells with angles between 60° and 90° were classified as perpendicular and those with angles between 0° and 30° were regarded as parallel. Cells orientated between 30° and 60° were considered non-classified.

### 3. Western blot and IP analyses

Analyses were performed as we described previously (65). For Western blotting, cells or tissues were lysed using 1.5x Laemmli buffer. Protein concentration was determined using the DC protein assay kit (Biorad). Equal amounts of proteins (10-30ug) were separated via SDS-PAGE. Proteins were transferred to a PVDF membrane (Biorad). The membrane was blocked with 10% BSA in TBST for 1 hour and incubated with a primary antibody at 4°C overnight. On the next day, the membrane was incubated with a Horseradish Peroxidase conjugated secondary antibody (ThermoFisher). The signals were detected using the Immobilon ECL Ultra Western HRP Substrate (Millipore) following instructions provided by the vendor. For co-IP analyses, tissues were lysed with lysis buffer (50mM Tris-HCl (pH 7.4), 0.1% Triton-100, 0.1% IGEPAL, 100mM NaCl, 10% glycine). Lysates were first incubated with an anti-GABPB antibody or an anti-SIRT7 antibody for 6 hours at 4°C and were then added with 30ul Protein G Magnet beads (Pierce) for incubation at 4°C overnight. On the next day, beads were thoroughly washed with lysis buffer and proteins were eluted by incubation with 1.5x Laemmli buffer at 70°C for 5 minutes. Supernatants were subjected to West analysis using proper antibodies. To test the impact of blocking AMPK on expression phosphorylated SIRT7, E13.5 embryonic hearts were isolated and cultured in DMEM (with 10% BSA) containing DMSO or 10uM Compound C (Millipore, #171620) or DMSO for 6 hours in a cell culture incubator (37 °C, 5% CO_2_). The RVs were then removed and were subjected to IP using an anti-SIRT7 antibody. The primary antibodies used for Western and IP analyses were purchased from Abcam (GABPA, ab224325; Phosphoserine, ab9332), Abclonal (RPS28, #A17937), BD Biosciences (N-Cadherin, #610920), Cell Signaling (Acetylated-Lysine, #9441; AMPKα, #2532; Phospho-AMPKα, 40H9; SIRT7, D3K5A; SP1, D4C3; YY1, D5D9Z), Iowa Hybridoma Bank (beta-Tubulin, E7), Proteintech (RPL35, #14826-1-AP; RPL37A, #14660-1-AP; RPS21, #16946-1-AP), and Santa Cruz (GABPB, E-7).

### 4. Culturing embryonic cardiomyocytes and measuring protein synthesis

Cardiomyocytes from RVs of E13.5 embryos were isolated using the neonatal heart dissociation kit purchased from Miltenyi Biotech. To help remove fibroblast contamination, isolated cells were pre-plated on a 100mm petri dish for 4 times in a cell culture incubator for 30 minutes. Remaining cells were then suspended in culture medium [DMEM with 10% FBS and 1xPen/Step (ThermoFisher)], plated in a 48-well culture plate, and cultured for 48 hours in cell culture incubator followed by subsequent procedures. De novo protein synthesis was measured using the Global Protein Synthesis Assay Kit (Abcam, ab235634) following the instruction provided by the vendor. Briefly, cells were incubated with 1x Protein Label (containing O-Propargyl-puromycin) added to the culture medium for 4 hours followed by signal detection. Total nuclei were visualized by DAPI staining. In a negative control experiment, cardiomyocytes isolated control embryos were treated with Cycloheximide (20ug/ml) for 30 minutes before adding Protein Label. Images were taken using AxioCam HRc high resolution digital camera connected to the Zeiss Axio fluorescent microscope under exactly identical settings between control and mutant samples. Signal intensity was quantified using Photoshop 11. To test the effect of blocking the AMPK activity, cultured cardiomyocytes were treated with 10uM Compound C for 16 hours followed by measuring protein synthesis.

### 5. TEM

E13.5 embryonic hearts were isolated from control and mutant embryos, treated with Ca++ free HEPES buffer (130mM NaCl, 5.4mM KCl, 0.33mM NaH_2_PO_4_, 0.5mM MgCl_2_, 22mM Glucose, and 20mM HEPES, pH 7.4) for 3 minutes, and then fixed with 2% paraformaldehyde/2.5% glutaraldehyde (EM grade, Sigma) for 24 hours at room temperature. Samples were then submitted to UAB High Resolution Imaging Facility for osmification, dehydration and embedding following the procedures described previously (67). Embedded samples were cut at 100nm and observed using the Tecnai Spirit T12 Transmission Electron Microscope (Thermo-Fisher, formerly FEI). Images were taken using an AMT (Advanced Microscopy Techniques, Corp) Bio Sprint 29 megapixel digital camera.

### 6. Measuring mitochondrial oxygen consumption and RCR

The RVs of control and cKO at E14.5 were dissected out and weighed using an AB54-S/FACT Analytical scale (Toledo). We usually combined 2-3 cKO RVs into one tube as the weight of a cKO RV was about half of a control RV (~1.2mg). Tissues were permeabilized as previously described (68). Briefly, samples were transferred to ice cold transport buffer (B1-200mM sucrose, 0.5mM Na2EDTA, 10mM Tris, 1g/L BSA) and were immediately submitted to the UAB Bioanalytical Redox Biology (BARB) Core. Tissues were permeabilized by gentle dissection on ice with needle tip forceps, washed in MiR03 buffer (0.5mM EGTA, 3mM MgCl_2_.6H2O, 20mM Taurine, 10mM KH_2_PO_4_, 20mM HEPES, 200mM Sucrose, 1g/L BSA) by gentle rotation for 15 minutes at 4°C, and then transferred to fresh MiR03 buffer for high resolution respirometry (HRR). HRR was performed by measuring oxygen consumption in 2mL of MiR03 buffer, in a two chamber respirometer (Oroboros Oxygraph-2k with DatLab software; Oroboros Instruments Corp., Innsbruck Austria) with constant stirring at 750 rpm. Respiration rates were measured using the protocols as previously described (15). Substrate mediated respiration (State 2 respiration or V_o_) was measured in the presence of complex I substrates (3mM Malate and 5mM Glutamate), followed by 1mM ADP (State 3 or V_max_). The respiratory control ratio (RCR) V_max_/V_o_ (State 3/State 2) was calculated using these measures. To evaluate inner mitochondrial membrane coupling, 10-100 µM ATR (added in 10 µM increments) was added to block ATP translocase and the oxygen consumption rate was then measured to obtain V_max_+ATR.

### 7. Measure the relative mitochondrial DNA copy numbers and ATP concentration

The relative mitochondrial DNA to nuclear DNA ratio was measured as described previously (69, 70) Total DNA was isolated from RVs of control and cKO hearts (E12.5) using the QIAamp DNA mini kit (Qiagen). qPCR analysis was performed in a Roche LightCycler 480 using primers specific for mitochondrial DNA (mMitoF1: 5’CTAGAAACCCCGAAACCAAA and mMitoR1: 5’ CCAGCTATCACCAAGCTCGT) and for nuclear DNA (mB2MF1: ATGGGAAGCCGAACATACTG and mB2MR1: 5’ CAGTCTCAGTGGGGGTGAAT). To measure ATP content, RVs and LVs were isolated from control and cKO embryos at E13.5 and weighed, followed by measuring ATP content using the Luminescent ATP Detection Assay kit (Abcam Ab113849). Luciferase activity was using the Synergy H1 microplate reader (Biotek). The raw numbers were normalized against tissue weight.

### 8. ScRNA-Seq and qRT-PCR

For scRNA-Seq, RVs of control and mutant embryos at E13.5 were isolated and treated with Trypsin to acquire single cells. Samples were pooled from 3-4 hearts of the same genotype. Cells were stained with 7-amino-actinomycin (7-AAD, ThermoFisher) to label nonviable cells and then sorted using a FACS Aria II sorter (BD BioSciences) to acquire viable cells. Single cell transcriptome libraries were prepared using 10xGenomics Single Cell 3ʹ v3 Reagent Kits according to the standard protocol outlined in the manual. The single cell libraries were sequenced using Illumina Next Seq500 with the goal of a minimum of 20,000 total reads per cell. The 10X Genomics Cellranger software (version 3.0.0), ‘mkfastq’, was used to create the fastq files from the sequencer. Following fastq file generation, Cellranger ‘count’ was used to align the raw sequence reads to the reference genome using STAR. The ‘count’ software created 3 data files (barcodes.tsv.gz, genes.tsv.gz, matrix.mtx.gz) from the ‘filtered_feature_bc_matrix’ folder that were loaded into the R (version 3.6.0) package Seurat version 3.0.2 (1,2), which allows for selection and filtration of cells based on QC metrics, data normalization and scaling, and detection of highly variable genes. We followed the two Seurat vignettes (https://satijalab.org/seurat/v3.0/pbmc3k_tutorial.html and https://satijalab.org/seurat/v3.0/immune_alignment.html) to create the Seurat data matrix object. In brief, we kept all genes expressed in ≥3 cells and cells with at least 200 detected genes. Cells with mitochondrial gene percentages over 7.5% and unique gene counts >5,000 or <200 were discarded as well. The data was normalized using Seurat’s ‘NormalizeData’ function, which uses a global-scaling normalization method, LogNormalize, to normalize the gene expression measurements for each cell to the total gene expression. The result is multiplied by a scale factor of 1e4 and the result is log-transformed. Highly variable genes were then identified using the function ‘FindVariableGenes’ in Seurat, which returned 2,000 features per dataset. We then merged the two datasets by identifying anchors using ‘FindIntegrationAnchors’ in Seurat using 30 dimensions followed by integrating the two datasets with ‘IntegrateData’. Following integration, we regressed out the variation arising from library size and percentage of mitochondrial genes using the function ‘ScaleData’ in Seurat. A UMAP was created to visualize the clusters of the combined datasets. We next identified conserved cell type markers with ‘FindConservedMarkers’ for each cluster. To identify differentially expressed genes in cell cluster across the two datasets, we used the function ‘FindMarkers’ in Seurat.

For qRT-PCR analysis, total RNA was isolated from RVs of control and mutant embryos at E13.5. Total RNA was reverse transcribed to acquire cDNA followed by real time PCR as we described previously (65). *Hprt* was used as the loading control. The primers for qRT-PCR were provided in Sup. Table 3.

### 9. Study approval

All procedures for mouse studies were approved by the Institutional Animal Care and Use Committee in UAB.

## Author contributions

Q. Z., S.Y., J. L., D. P., H. W., and S. L. performed the experiments and acquired data. D. C. and S. L. analyzed data. Q. Z., S. Y., Q. W., K. M., K. L., and K. J. designed the research studies, analyzed data and wrote the manuscript. H. S. provided the *Drp1^loxp/loxp^* mouse line and helped write the manuscript.

## Acknowledgments

We thank Dr. G. A. Porter (University of Rochester Medical Ctr.) for suggestions on isolating and culturing embryonic cardiomyocytes, Dr. M. Wu (Univ. of Houston) for suggestions on measuring cardiomyocyte orientation, Dr. S. Kubalak (Medical University of South Carolina) for providing the MLC2A antibody, and Dr. S. H. Litovsky (UAB) for helping with analyzing TEM images. We thank UAB Genomics Core Facility for performing deep-Seq and Sanger Sequencing, E. Phillips and M. Foley at UAB High Resolution Imaging Facility for performing TEM studies, and K. Smith-Johnston and M. J. Sammy at the UAB Bioanalytical Redox Biology (BARB) Core for performing HRR. We thank members in Dr. Jiao’s lab for suggestions on this project. This study was supported by a grant from NIH (R01HL095783) and a UAB internal AMC21 grant awarded to K. J. UAB BARB Core is supported by NIDDK DK P30DK079626), NORC (NIDDK DK056336), UAB Center for Exercise Medicine (UCEM), Comprehensive Diabetes Center (UCDC), Center for Free Radical Biology (CFRB) and Comprehensive Neuroscience Center (UCNC).

---

**Sup. Fig. 1.**
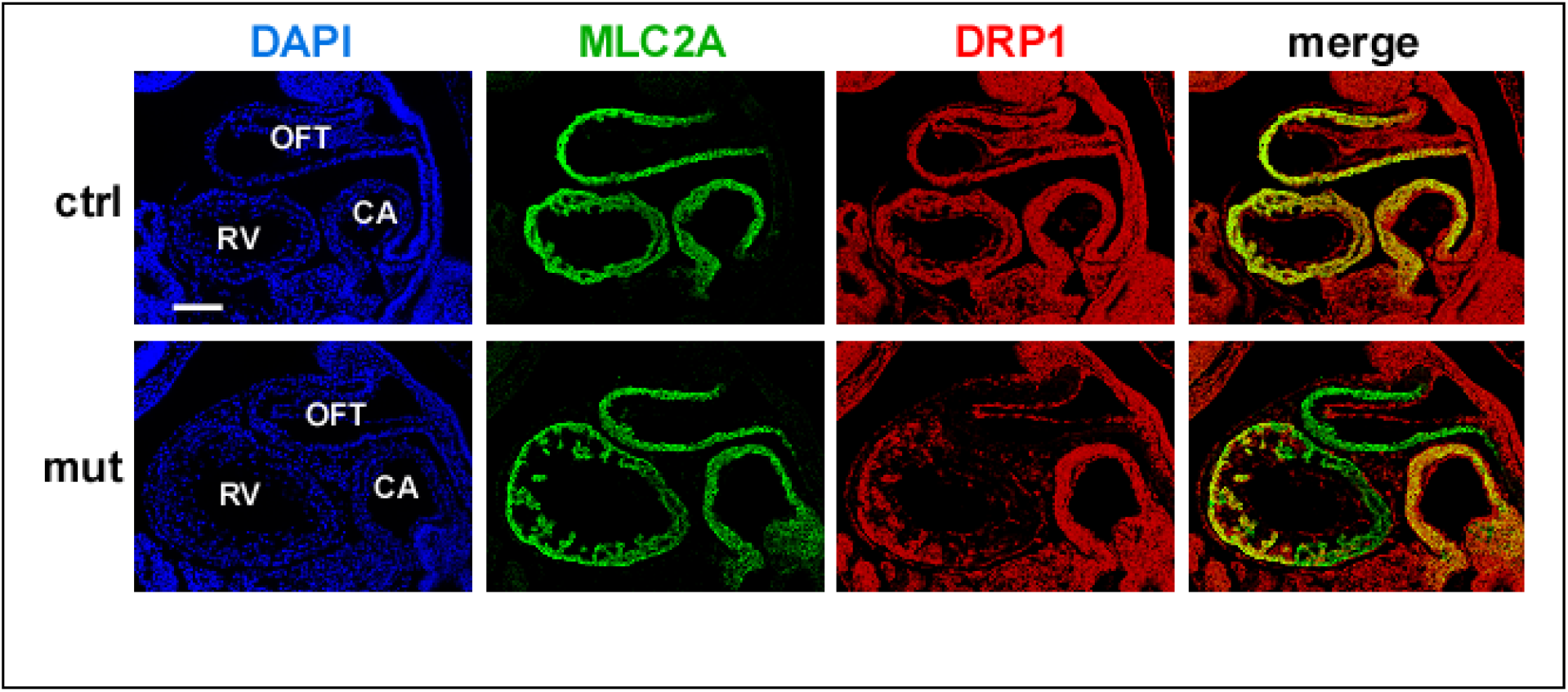
Expression of DRP1 in E9.5 embryos. *Mef2c-AHF-Cre/Drp1^loxp/+^* male mice were crossed with *Drp1^loxp/loxp^* female mice to obtain mutant (Mut, *Mef2c-AHF-Cre/Drp1^loxp/loxp^*) and control (Ctrl, *Drp1^loxp/+^* or *Drp1^loxp/loxp^*) embryos at E9.5. Sagittal sections were immunostained with antibodies against DRP1 (red) and MLC1A (red, a cardiomyocyte marker). The outflow tract (OFT), atrial ventricular canal (AVC) and common atrium (CA) regions are shown. The overt red signal (indicating DRP1 expression) was reduced in the OFT and the half of the RV that was adjacent to the OFT, while remained strong in the other half of the RV.

**Sup. Fig. 2.**
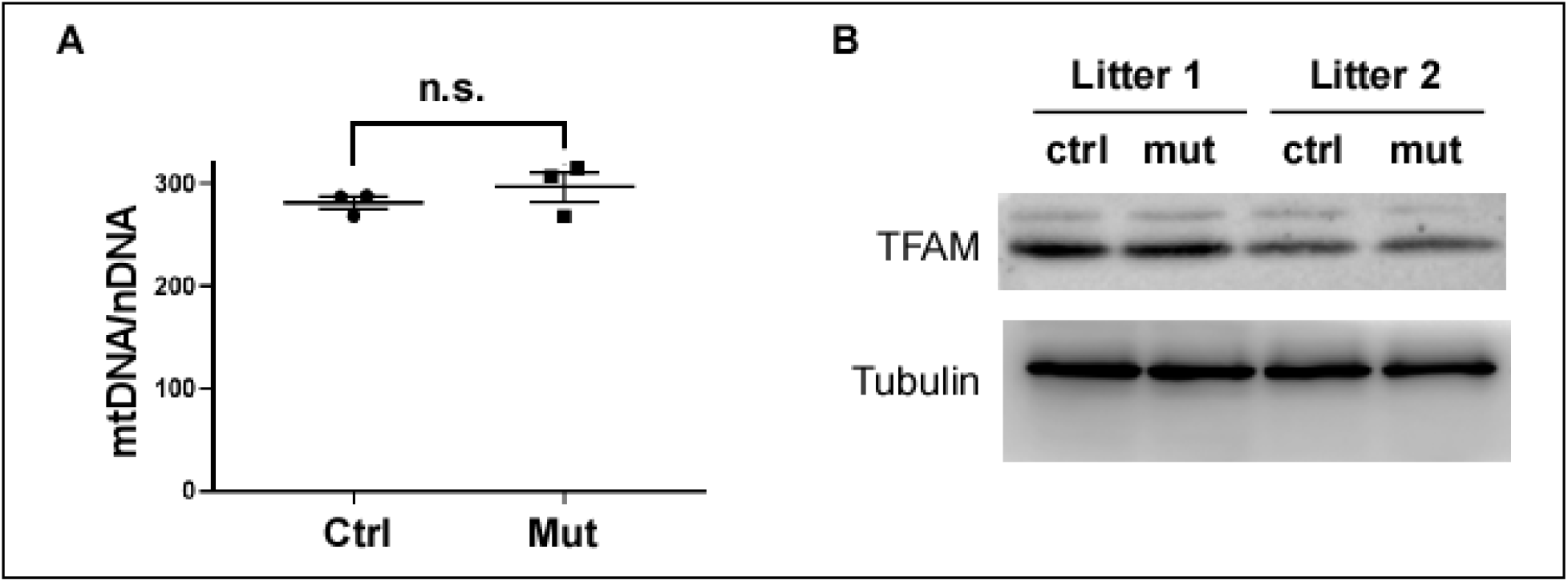
General mitochondrial biogenesis was not impaired by *Drp1*. Deletion. **(A)** The relative mitochondrial DNA to nuclear DNA ratio (mtDNA/nDNA) in control and mutant RVs at E12.5 was determined as described in Methods. n.s.: not significant, two tailed unpaired Student’s t test. **(B)** Western analysis was performed using protein lysates isolated from control and mutant RVs. Tubulin was used as the loading control.

**Sup. Fig. 3.**
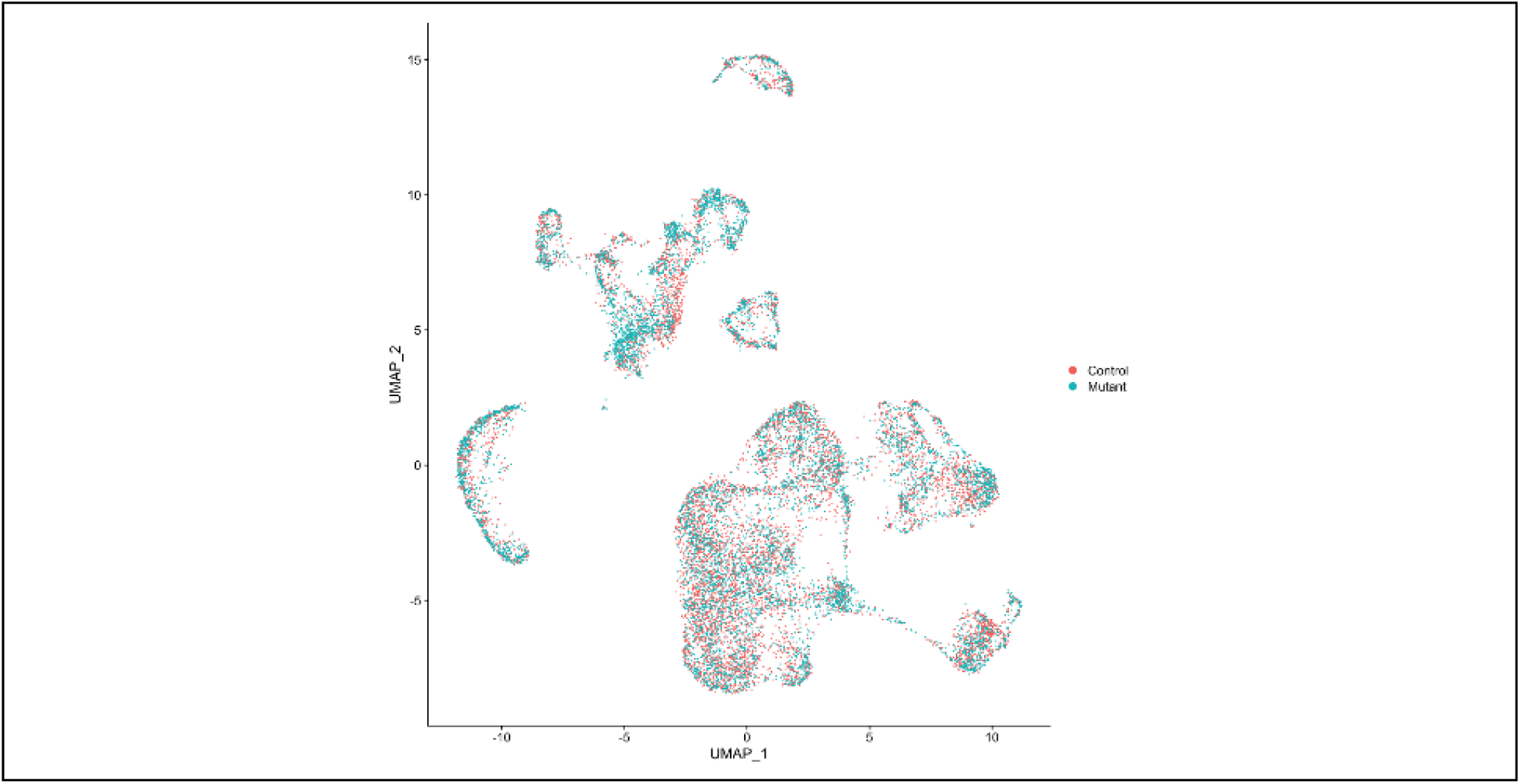
**(A)** UMAP chart showing the distribution of control cardiomyocytes (red) and mutant cardiomyocytes (blue) across all clusters.

**Sup. Fig. 4.**
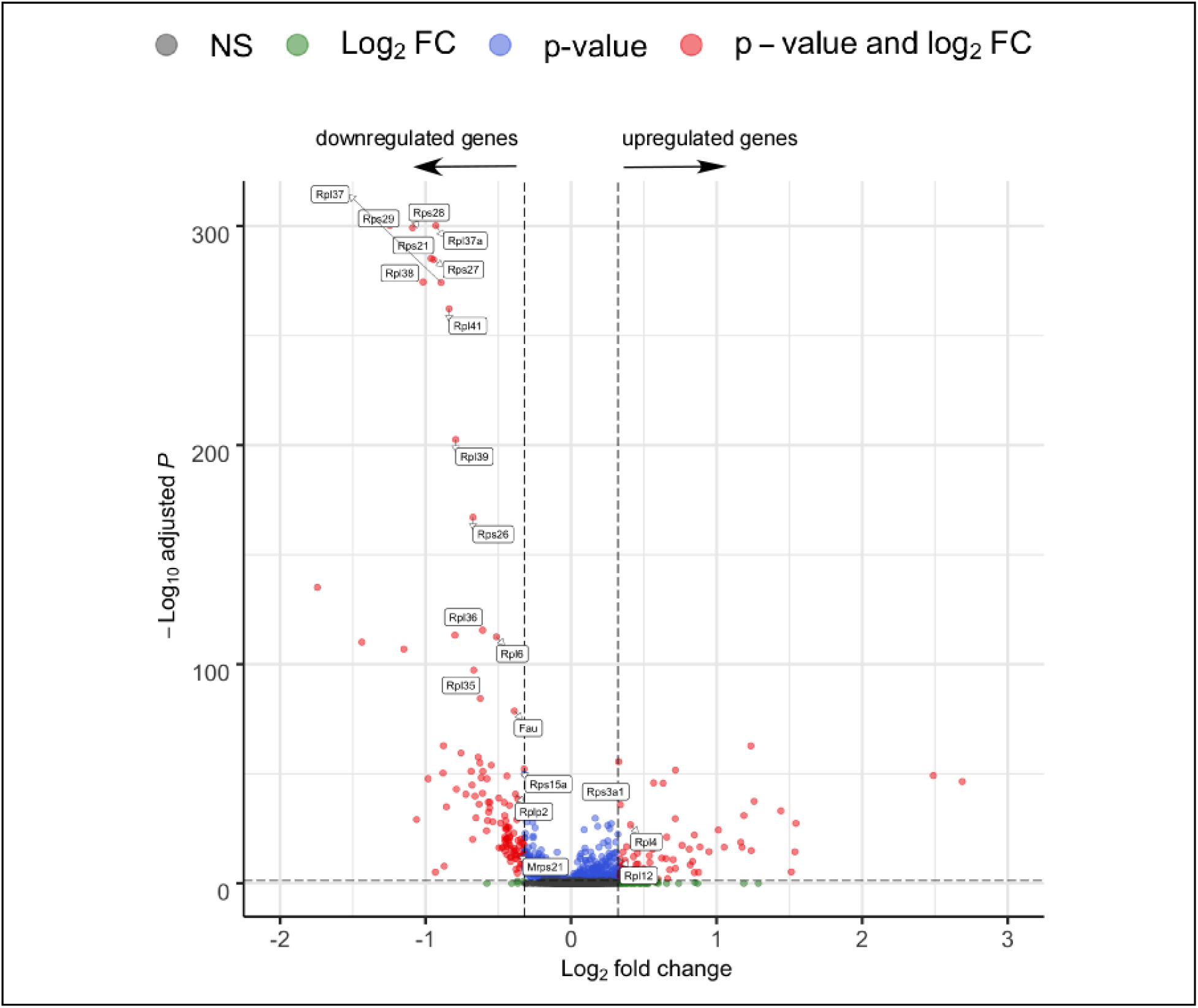
Enlarged volcano chart. (corresponding to Fig. 6C).

## Reference

1. Ernster L, and Schatz G. Mitochondria: a historical review. J Cell Biol. 1981;91(3 Pt 2):227s–55s.

2. Friedman JR, and Nunnari J. Mitochondrial form and function. Nature. 2014;505(7483):335–43.

3. Dorn GW, 2nd, Vega RB, and Kelly DP. Mitochondrial biogenesis and dynamics in the developing and diseased heart. Genes Dev. 2015;29(19):1981–91.

4. Dorn G, II. Mitochondrial fission/fusion and cardiomyopathy. Curr Opin Genet Dev. 2016;38:38–44.

5. Zhao Q, Sun Q, Zhou L, Liu K, and Jiao K. Complex Regulation of Mitochondrial Function During Cardiac Development. J Am Heart Assoc. 2019;8(13):e012731.

6. Sharma A, Smith HJ, Yao P, and Mair WB. Causal roles of mitochondrial dynamics in longevity and healthy aging. EMBO Rep. 2019;20(12):e48395.

7. Schiff M, Ogier de Baulny H, and Lombes A. Neonatal cardiomyopathies and metabolic crises due to oxidative phosphorylation defects. Semin Fetal Neonatal Med. 2011;16(4):216–21.

8. El-Hattab AW, and Scaglia F. Mitochondrial Cardiomyopathies. Front Cardiovasc Med. 2016;3:25.

9. Berardo A, Musumeci O, and Toscano A. Cardiological manifestations of mitochondrial respiratory chain disorders. Acta Myol. 2011;30(1):9–15.

10. Sylva M, van den Hoff MJ, and Moorman AF. Development of the human heart. American Journal of Medical Genetics Part A. 2014;164(6):1347–71.

11. Christoffels VM, Burch JB, and Moorman AF. Architectural plan for the heart: early patterning and delineation of the chambers and the nodes. Trends in cardiovascular medicine. 2004;14(8):301–7.

12. Buckingham M, Meilhac S, and Zaffran S. Building the mammalian heart from two sources of myocardial cells. Nat Rev Genet. 2005;6(11):826–35.

13. Mackler B, Grace R, and Duncan HM. Studies of mitochondrial development during embryogenesis in the rat. Archives of biochemistry and biophysics. 1971;144(2):603–10.

14. Hom JR, Quintanilla RA, Hoffman DL, de Mesy Bentley KL, Molkentin JD, Sheu SS, et al. The permeability transition pore controls cardiac mitochondrial maturation and myocyte differentiation. Dev Cell. 2011;21(3):469–78.

15. Beutner G, Eliseev RA, and Porter GA Jr. Initiation of electron transport chain activity in the embryonic heart coincides with the activation of mitochondrial complex 1 and the formation of supercomplexes. PLoS One. 2014;9(11):e113330.

16. Lopaschuk GD, and Jaswal JS. Energy metabolic phenotype of the cardiomyocyte during development, differentiation, and postnatal maturation. Journal of cardiovascular pharmacology. 2010;56(2):130–40.

17. Chung S, Dzeja PP, Faustino RS, Perez-Terzic C, Behfar A, and Terzic A. Mitochondrial oxidative metabolism is required for the cardiac differentiation of stem cells. Nature clinical practice Cardiovascular medicine. 2007;4 Suppl 1:S60–7.

18. Scarpulla RC. Transcriptional paradigms in mammalian mitochondrial biogenesis and function. Physiol Rev. 2008;88(2):611–38.

19. Tilokani L, Nagashima S, Paupe V, and Prudent J. Mitochondrial dynamics: overview of molecular mechanisms. Essays Biochem. 2018;62(3):341–60.

20. Kageyama Y, Zhang Z, and Sesaki H. Mitochondrial division: molecular machinery and physiological functions. Curr Opin Cell Biol. 2011;23(4):427–34.

21. Kasahara A, Cipolat S, Chen Y, Dorn GW, 2nd, and Scorrano L. Mitochondrial fusion directs cardiomyocyte differentiation via calcineurin and Notch signaling. Science. 2013;342(6159):734–7.

22. Chen Y, Liu Y, and Dorn GW, 2nd. Mitochondrial fusion is essential for organelle function and cardiac homeostasis. Circ Res. 2011;109(12):1327–31.

23. Chen H, Detmer SA, Ewald AJ, Griffin EE, Fraser SE, and Chan DC. Mitofusins Mfn1 and Mfn2 coordinately regulate mitochondrial fusion and are essential for embryonic development. J Cell Biol. 2003;160(2):189–200.

24. Detmer SA, and Chan DC. Complementation between mouse Mfn1 and Mfn2 protects mitochondrial fusion defects caused by CMT2A disease mutations. J Cell Biol. 2007;176(4):405–14.

25. Ishihara T, Ban-Ishihara R, Maeda M, Matsunaga Y, Ichimura A, Kyogoku S, et al. Dynamics of mitochondrial DNA nucleoids regulated by mitochondrial fission is essential for maintenance of homogeneously active mitochondria during neonatal heart development. Mol Cell Biol. 2015;35(1):211–23.

26. Kageyama Y, Hoshijima M, Seo K, Bedja D, Sysa-Shah P, Andrabi SA, et al. Parkin-independent mitophagy requires Drp1 and maintains the integrity of mammalian heart and brain. EMBO J. 2014;33(23):2798–813.

27. Verzi MP, McCulley DJ, De Val S, Dodou E, and Black BL. The right ventricle, outflow tract, and ventricular septum comprise a restricted expression domain within the secondary/anterior heart field. Dev Biol. 2005;287(1):134–45.

28. Guo X, Aviles G, Liu Y, Tian R, Unger BA, Lin YT, et al. Mitochondrial stress is relayed to the cytosol by an OMA1-DELE1-HRI pathway. Nature. 2020;579(7799):427–32.

29. Desai BN, Myers BR, and Schreiber SL. FKBP12-rapamycin-associated protein associates with mitochondria and senses osmotic stress via mitochondrial dysfunction. Proc Natl Acad Sci U S A. 2002;99(7):4319–24.

30. Dennis PB, Jaeschke A, Saitoh M, Fowler B, Kozma SC, and Thomas G. Mammalian TOR: a homeostatic ATP sensor. Science. 2001;294(5544):1102–5.

31. Topf U, Uszczynska-Ratajczak B, and Chacinska A. Mitochondrial stress-dependent regulation of cellular protein synthesis. J Cell Sci. 2019;132(8).

32. Wakabayashi J, Zhang Z, Wakabayashi N, Tamura Y, Fukaya M, Kensler TW, et al. The dynamin-related GTPase Drp1 is required for embryonic and brain development in mice. J Cell Biol. 2009;186(6):805–16.

33. Weisz SH, Limongelli G, Pacileo G, Calabro P, Russo MG, Calabro R, et al. Left ventricular non compaction in children. Congenital heart disease. 2010;5(5):384–97.

34. Towbin JA, and Jefferies JL. Cardiomyopathies Due to Left Ventricular Noncompaction, Mitochondrial and Storage Diseases, and Inborn Errors of Metabolism. Circ Res. 2017;121(7):838–54.

35. Miao L, Li J, Li J, Lu Y, Shieh D, Mazurkiewicz JE, et al. Cardiomyocyte orientation modulated by the Numb family proteins-N-cadherin axis is essential for ventricular wall morphogenesis. Proc Natl Acad Sci U S A. 2019;116(31):15560–9.

36. Li J, Miao L, Shieh D, Spiotto E, Li J, Zhou B, et al. Single-Cell Lineage Tracing Reveals that Oriented Cell Division Contributes to Trabecular Morphogenesis and Regional Specification. Cell Rep. 2016;15(1):158–70.

37. Baker MJ, Frazier AE, Gulbis JM, and Ryan MT. Mitochondrial protein-import machinery: correlating structure with function. Trends Cell Biol. 2007;17(9):456–64.

38. Mitra K, Rikhy R, Lilly M, and Lippincott-Schwartz J. DRP1-dependent mitochondrial fission initiates follicle cell differentiation during Drosophila oogenesis. J Cell Biol. 2012;197(4):487–97.

39. Song M, Mihara K, Chen Y, Scorrano L, and Dorn GW, 2nd. Mitochondrial fission and fusion factors reciprocally orchestrate mitophagic culling in mouse hearts and cultured fibroblasts. Cell metabolism. 2015;21(2):273–86.

40. Becht E, McInnes L, Healy J, Dutertre CA, Kwok IWH, Ng LG, et al. Dimensionality reduction for visualizing single-cell data using UMAP. Nature biotechnology. 2018.

41. Li G, Xu A, Sim S, Priest JR, Tian X, Khan T, et al. Transcriptomic Profiling Maps Anatomically Patterned Subpopulations among Single Embryonic Cardiac Cells. Dev Cell. 2016;39(4):491–507.

42. Hill MC, Kadow ZA, Li L, Tran TT, Wythe JD, and Martin JF. A cellular atlas of Pitx2-dependent cardiac development. Development. 2019;146(12).

43. Zhou Y, Zhou B, Pache L, Chang M, Khodabakhshi AH, Tanaseichuk O, et al. Metascape provides a biologist-oriented resource for the analysis of systems-level datasets. Nature communications. 2019;10(1):1523.

44. Perry RP. The architecture of mammalian ribosomal protein promoters. BMC Evol Biol. 2005;5:15.

45. Nosrati N, Kapoor NR, and Kumar V. Combinatorial action of transcription factors orchestrates cell cycle-dependent expression of the ribosomal protein genes and ribosome biogenesis. FEBS J. 2014;281(10):2339–52.

46. Li X, Zheng Y, Hu H, and Li X. Integrative analyses shed new light on human ribosomal protein gene regulation. Sci Rep. 2016;6:28619.

47. Rosmarin AG, Resendes KK, Yang Z, McMillan JN, and Fleming SL. GA-binding protein transcription factor: a review of GABP as an integrator of intracellular signaling and protein-protein interactions. Blood Cells Mol Dis. 2004;32(1):143–54.

48. Hoffmeyer A, Avots A, Flory E, Weber CK, Serfling E, and Rapp UR. The GABP-responsive element of the interleukin-2 enhancer is regulated by JNK/SAPK-activating pathways in T lymphocytes. J Biol Chem. 1998;273(17):10112–9.

49. Fromm L, and Burden SJ. Neuregulin-1-stimulated phosphorylation of GABP in skeletal muscle cells. Biochemistry. 2001;40(17):5306–12.

50. Ryu D, Jo YS, Lo Sasso G, Stein S, Zhang H, Perino A, et al. A SIRT7-dependent acetylation switch of GABPbeta1 controls mitochondrial function. Cell metabolism. 2014;20(5):856–69.

51. Sun L, Fan G, Shan P, Qiu X, Dong S, Liao L, et al. Regulation of energy homeostasis by the ubiquitin-independent REGgamma proteasome. Nature communications. 2016;7:12497.

52. Parker DJ, Iyer A, Shah S, Moran A, Hjelmeland AB, Basu MK, et al. A new mitochondrial pool of cyclin E, regulated by Drp1, is linked to cell-density-dependent cell proliferation. J Cell Sci. 2015;128(22):4171–82.

53. Kashatus JA, Nascimento A, Myers LJ, Sher A, Byrne FL, Hoehn KL, et al. Erk2 phosphorylation of Drp1 promotes mitochondrial fission and MAPK-driven tumor growth. Mol Cell. 2015;57(3):537–51.

54. Serasinghe MN, Wieder SY, Renault TT, Elkholi R, Asciolla JJ, Yao JL, et al. Mitochondrial division is requisite to RAS-induced transformation and targeted by oncogenic MAPK pathway inhibitors. Mol Cell. 2015;57(3):521–36.

55. Simula L, Pacella I, Colamatteo A, Procaccini C, Cancila V, Bordi M, et al. Drp1 Controls Effective T Cell Immune-Surveillance by Regulating T Cell Migration, Proliferation, and cMyc-Dependent Metabolic Reprogramming. Cell Rep. 2018;25(11):3059–73 e10.

56. Vantaggiato C, Castelli M, Giovarelli M, Orso G, Bassi MT, Clementi E, et al. The Fine Tuning of Drp1-Dependent Mitochondrial Remodeling and Autophagy Controls Neuronal Differentiation. Front Cell Neurosci. 2019;13:120.

57. Wu MJ, Chen YS, Kim MR, Chang CC, Gampala S, Zhang Y, et al. Epithelial-Mesenchymal Transition Directs Stem Cell Polarity via Regulation of Mitofusin. Cell metabolism. 2019;29(4):993–1002 e6.

58. Wu S, and Zou MH. AMPK, Mitochondrial Function, and Cardiovascular Disease. Int J Mol Sci. 2020;21(14).

59. Lindqvist LM, Tandoc K, Topisirovic I, and Furic L. Cross-talk between protein synthesis, energy metabolism and autophagy in cancer. Curr Opin Genet Dev. 2018;48:104–11.

60. Quiros PM, Prado MA, Zamboni N, D’Amico D, Williams RW, Finley D, et al. Multi-omics analysis identifies ATF4 as a key regulator of the mitochondrial stress response in mammals. J Cell Biol. 2017;216(7):2027–45.

61. Tanwar DK, Parker DJ, Gupta P, Spurlock B, Alvarez RD, Basu MK, et al. Crosstalk between the mitochondrial fission protein, Drp1, and the cell cycle is identified across various cancer types and can impact survival of epithelial ovarian cancer patients. Oncotarget. 2016;7(37):60021–37.

62. Liang R, Arif T, Kalmykova S, Kasianov A, Lin M, Menon V, et al. Restraining Lysosomal Activity Preserves Hematopoietic Stem Cell Quiescence and Potency. Cell stem cell. 2020;26(3):359–76.e7.

63. Peng Y, Song L, Li D, Kesterson R, Wang J, Wang L, et al. Sema6D acts downstream of bone morphogenetic protein signalling to promote atrioventricular cushion development in mice. Cardiovasc Res. 2016;112(2):532–42.

64. Liu Y, Harmelink C, Peng Y, Chen Y, Wang Q, and Jiao K. CHD7 interacts with BMP R-SMADs to epigenetically regulate cardiogenesis in mice. Hum Mol Genet. 2014;23(8):2145–56.

65. Yan S, Thienthanasit R, Chen D, Engelen E, Bruhl J, Crossman DK, et al. CHD7 regulates cardiovascular development through ATP-dependent and -independent activities. Proc Natl Acad Sci U S A. 2020;117(46):28847–58.

66. Harmelink C, Peng Y, Debenedittis P, Chen H, Shou W, and Jiao K. Myocardial Mycn is essential for mouse ventricular wall morphogenesis. Dev Biol. 2013;373(1):53–63.

67. Larson-Casey JL, Deshane JS, Ryan AJ, Thannickal VJ, and Carter AB. Macrophage Akt1 Kinase-Mediated Mitophagy Modulates Apoptosis Resistance and Pulmonary Fibrosis. Immunity. 2016;44(3):582–96.

68. Hunter GR, Moellering DR, Carter SJ, Gower BA, Bamman MM, Hornbuckle LM, et al. Potential Causes of Elevated REE after High-Intensity Exercise. Med Sci Sports Exerc. 2017;49(12):2414–21.

69. Singh B, Li X, Owens KM, Vanniarajan A, Liang P, and Singh KK. Human REV3 DNA Polymerase Zeta Localizes to Mitochondria and Protects the Mitochondrial Genome. PLoS One. 2015;10(10):e0140409.

70. Quiros PM, Goyal A, Jha P, and Auwerx J. Analysis of mtDNA/nDNA Ratio in Mice. Curr Protoc Mouse Biol. 2017;7(1):47–54.

